# Natural variation in physical responses to waterlogging across climate-diverse pennycress accessions

**DOI:** 10.1101/2024.08.20.608872

**Authors:** Rachel Combs-Giroir, John Lagergren, Daniel A. Jacobson, Andrea R. Gschwend

## Abstract

Fluctuations in flooding and differences in hydrological gradients affect the geographic distribution of plant species across ecosystems, resulting in the presence of adaptive traits in populations that confer enhanced fitness in flooded environments. Many winter annual crops, such as pennycress (*Thlaspi arvense* L.), are subjected to heavy precipitation events during their growing season, which are increasing in frequency due to climate change. Therefore, it is essential to identify pennycress accessions with natural variation in flooding resilience. In this study, we used climate modeling data to assess spring soil moisture levels in the geographic origins of 471 natural pennycress accessions. We selected 34 accessions with variations in predicted soil moisture to test survivability under prolonged waterlogging at the rosette stage. This identified variation in waterlogging tolerance, where six accessions had 0% survivability and nine accessions had 100% survivability. It took at least seven weeks for the first accessions to die under waterlogging, indicating that pennycress is hardy to prolonged waterlogging at the vegetative stage. Furthermore, we chose three “susceptible” and five “tolerant” accessions to waterlog for one week at the reproductive stage, the growth stage aligned with heavy spring rainfall. Six accessions had significantly reduced seed weight at maturity after recovery from waterlogging, and two accessions had minimal impacts on growth and seed yield after waterlogging. These two accessions can be used in future studies to explore adaptive traits, such as changes in root characteristics, as well as the genetic variation that contributes to pennycress waterlogging tolerance.

## Introduction

As a result of global climate change, crops are increasingly exposed to fluctuating water availability, such as severe drought or various types of flooding stress (Anderson et al. 2020; Watanabe et al. 2018). Flooding stress can occur in the form of waterlogging, where the root system is fully saturated, or in the form of submergence, where above-ground tissues are partially or fully submerged under water (Sasidharan et al. 2017). This creates a disastrous effect on economically important crops, because flooding stress reduces plant growth and yield, leading to an estimated annual loss of $74 billion globally (Kaur et al. 2020). One reason for reduced plant growth under flooding is a slow diffusion of oxygen in water as opposed to air, which exposes flooded tissues to low rates of oxidative phosphorylation and ATP production, ultimately putting the plant in a severe energy crisis (Bailey-Serres et al. 2012). Other causes for impaired growth and development of flooded plants are decreased nutrient uptake and photosynthetic activity (Elzenga and van Veen 2010; Mommer et al. 2006; Wollmer, Pitann and K.-H. Mühling 2018), as well as increased amounts of reactive oxygen species and oxidative stress (León et al. 2020; Liu and Zwiazek 2022).

Variations in hydrological gradients and flooding frequencies across ecosystems influence the natural geographic distribution of plant species (Voesenek et al. 2004). These variations in ecosystems lead to selection pressures for flooding response adaptive traits and survival strategies, such as the quiescence and escape strategies, adventitious root production, and aerenchyma formation (Bailey- Serres and Voesenek 2008; Colmer and Voesenek 2009; Combs-Giroir and Gschwend 2024). Plants that utilize the quiescence strategy limit growth and conserve substrates until the floodwaters recede, whereas plants demonstrating the escape strategy initiate shoot elongation to emerge from the floodwaters, which facilitates oxygen transport to hypoxic tissues (Akman et al. 2012; Bailey-Serres and Voesenek 2008; Colmer and Voesenek 2009). Gas exchange is also facilitated by adventitious roots that emerge on the stem or hypocotyl (Steffens and Rasmussen 2016) and root aerenchyma, or longitudinally connected gas spaces (Armstrong 1980). These adaptive traits have enabled different plant species or populations to survive various flooding conditions.

Pennycress, *Thlaspi arvense* L., is a winter annual oilseed and a promising new biofuel cash crop for integration into corn and soybean rotations across the U.S. Midwest (McGinn et al. 2019; Phippen et al. 2022; Sedbrook et al. 2014). Pennycress not only provides ecological benefits over the winter to otherwise fallow fields, it also produces seed oil that can be turned into renewable diesel and aviation fuel, and seed meal that can be used for high protein animal feed (Phippen et al. 2022). However, pennycress fields are exposed to many different abiotic threats throughout the spring, when plants are flowering and producing seeds, including heavy precipitation. Previously, Combs-Giroir et al. studied the impact of one week of waterlogging during the reproductive stage on two pennycress accessions, which revealed negative impacts on yield and biomass, as well as the reconfiguration of energy metabolism through upregulation of genes involved in fermentation and glycolysis (Combs-Giroir et al. 2024). Pennycress is native to Eurasia, but can be found throughout temperate regions worldwide (Geng et al. 2021; Holm et al. 1997; Warwick et al. 2002). Given its wide distribution, some natural pennycress populations have likely evolved to survive in the various climatic conditions of their local habitats, including those with regularly wet or waterlogged soil, possibly through similar adaptive traits observed in other *Brassicaceae* species (Combs-Giroir and Gschwend 2024). We predict that these accessions would be waterlogging tolerant, meaning they would be able to survive prolonged waterlogging with minimal impacts on seed yield and biomass.

A germplasm bank containing 471 globally collected natural pennycress accessions, mostly from North America and Europe, has been curated by the Integrated Pennycress Resilience Project (IPReP) (Sedbrook et al., in preparation). These accessions have been collected from various habitats. Pennycress is a ruderal species and is most commonly found where the soil has been disturbed, for example along the edges of farm fields, marshy regions of fields, areas of construction, including roadwork, and where animals disturb the soil. In this study, we utilized climate modeling data to select 34 of these natural pennycress accessions that were collected from geographic locations with varying predicted soil water availability and screened them for survivability under waterlogging. We found variation in waterlogging tolerance among these 34 accessions under prolonged waterlogging imposed at the rosette stage. We further selected 8 of these accessions with low and high waterlogging tolerance to test their performance under waterlogging at the reproductive stage, the growth stage that aligns with the month of April, when heavy precipitation is common. This revealed significant variation in waterlogging tolerance among the accessions tested and led to the identification of two waterlogging-tolerant pennycress accessions. This knowledge will accelerate the discovery of physical and molecular waterlogging stress adaptations in wild pennycress populations, which can be used for introgression into high-yielding pennycress lines through breeding programs.

## Materials/Methods

### Plant Material

471 pennycress natural accession source seeds were collected from various locations, mostly across North America and Europe, by citizens, Integrated Pennycress Resilience Project (IPReP) collaborators, or other research teams and were made available for this project by the Germplasm Resource Information Network (GRIN), Arabidopsis Biological Resource Center (ABRC), and Illinois State University (ISU; John Sedbrook) (Galanti et al. 2022; Nunn et al. 2022; USDA ARS 2024; ABRC n.d.; Sedbrook et al., in preparation) (Figure 1A; Supplemental Table 1). The accessions were grown to increase seed number by IPReP collaborators (Western Illinois University, Illinois State University, University of Minnesota, Washington State University, Donald Danforth Plant Sciences Laboratory, Carnegie Institution for Science, and The Ohio State University), and a subset were then used for our waterlogging experiments. Accessions ISU_381 and ISU_378 were collected by Patrick Bulman and accessions OSU_1, OSU_2, OSU_5, and OSU_6 were collected by Rachel Combs-Giroir from known wet locations such as ditches, waterlogged fields, marshy areas, or riverbanks, and original source seed for these accessions were used for the rosette stage waterlogging experiments. MN106 and Spring 32-10 are reference lines and were obtained from Illinois State University and Western Illinois University and seed increases were conducted at OSU.

**Figure 1.**
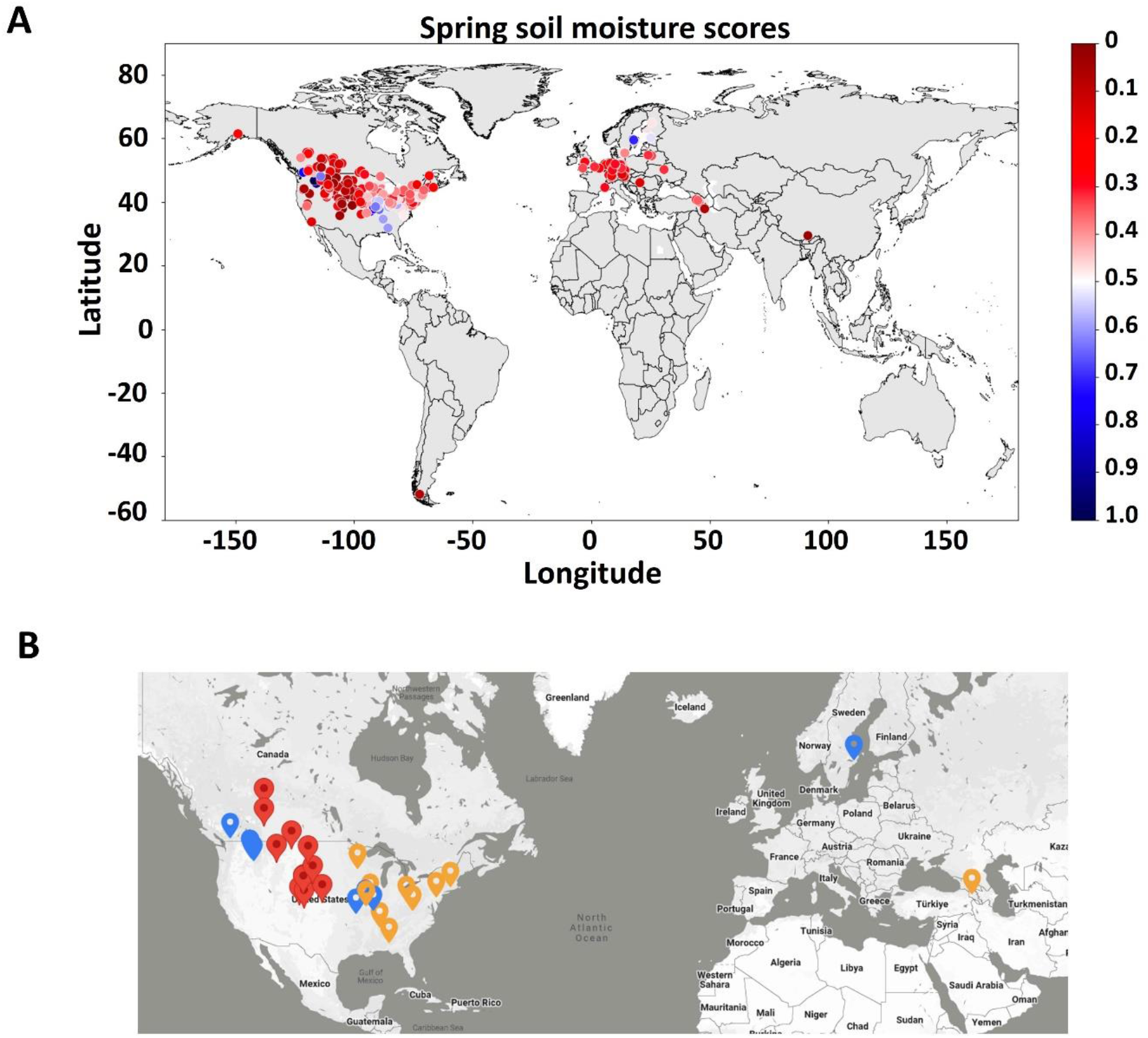
A) Spring soil water availability scores for 471 natural pennycress accessions based on soil moisture climate data for their geographic origins during March, April, and May between 2009 and 2018. Blue dots correspond to areas with high soil water availability (predicted wetter environments) and red dots correspond to areas with low soil water availability (predicted drier environments). The bar graph represents the distribution of the soil water availability on a scale of 0 to 1, with lower scores or drier climates on the left. B) A map of the source population locations for the 34 pennycress natural accessions used for rosette-stage screening under waterlogging, selected based on spring soil water availability scores. Red pins indicate accessions with a soil water availability score of 0-0.25, orange pins indicate accessions with a soil water availability score of 0.25-0.65, and blue pins indicate a soil water availability score of 0.65-1.

### Modeling Effective Water Availability

A soil water availability (SWA) score was developed to identify climate divergence among 471 natural pennycress accessions. Climate data was pulled from the TerraClimate repository (Abatzoglou et al. 2018) based on the closest location to the GPS coordinates of each pennycress natural accession. The closest point was computed by minimizing the Euclidean distance in latitude/longitude coordinates. The effective water availability, or the SWA score, was predicted based on soil moisture which is calculated by subtracting actual evapotranspiration and runoff from total precipitation/rain accumulation. Spring SWA was computed by taking the average of all soil moisture observations from March, April, and May between 2009 and 2018. Each observation was normalized by the minimum and maximum value to produce a score that ranged from 0 to 1. Low scores indicated dry locations with low soil moisture and high scores indicated wet locations with high soil moisture. All scripts were created using Python (version 3.8.10) and its libraries (https://github.com/jlager/pennycress-soil-moisture). These SWA scores were projected onto a map to reveal geospatial trends related to waterlogging (Figure 1A).

### Growth Conditions

Seeds were sterilized with a 70% ethanol rinse, followed by 10-minute incubation in 30% bleach/0.01% SDS solution, and then rinsed 3 times with sterile water. Seeds were then plated on petri dishes between two pieces of Whatman paper with 2mL sterile water, stratified at 4°C for 5 days, and moved to a growth chamber at 21°C with a photoperiod of 16-hour light/8-hour dark. Following 1 week of germination, all the seedlings were vernalized at 4°C under 14hrs of daily light for 3 weeks. After vernalization, the seedlings were transplanted into plug cells in Berger BM2 Germination Mix (Berger, Saint-Modeste, QC, Canada) and then into 4-inch pots after the first true leaves fully developed. The plants were grown at ∼21°C with periods of 16-hour light/8-hour dark either in a greenhouse (rosette stage experiment) or growth chamber (reproductive stage experiment). For the rosette stage experiment, the media in the pots was an even ratio of PRO-MIX BX MYCORRHIZAE (Premier Horticulture, Quakertown, PA, United States) and Miracle-Gro All Purpose Potting mix (Scotts Miracle-Gro, Marysville, OH, United States). For the reproductive stage experiment, the media in the pots was an even ratio of Turface MVP calcined clay (PROFILE Products LLC, Buffalo Grove, IL United States), PRO-MIX BX MYCORRHIZAE (Premier Horticulture, Quakertown, PA, United States), and Miracle-Gro All Purpose Potting mix (Scotts Miracle-Gro, Marysville, OH, United States). Plants in the reproductive stage experiment were watered with Jack’s Nutrients 12-4- 16 RO (jacksnutrients.com) at 100 ppm 2-3 times a week.

### Experimental Design – Rosette Stage

Thirty-four accessions (Supplemental Table 1 and Figure 1B) were selected for survivability screening under prolonged waterlogging at the rosette stage. When plants were 50 days old, four plants per accession were randomly placed inside 7.6-liter buckets (three plants of different accessions per bucket) and two plants per accession were randomly placed in 10” x 20” trays (10 plants total per tray) in a completely randomized design. The plants in the buckets were then waterlogged by filling the buckets with tap water up to the soil line at the top of the pots (treatment). The plants in the trays were bottom-watered normally as controls approximately every other day so that the soil was completely saturated after watering, but dry before the next watering. Waterlogged plants were continuously waterlogged until they died or fully senesced and the water levels were maintained daily. The plants in the trays were grown to maturity, or full senescence, under a normal watering regime. Survivability was measured by the number of days it took until the plant was dead or had fully senesced, regardless of the stage of development. Total seed weight was measured for plants that flowered and produced seed.

### Experimental Design – Reproductive Stage

Eight accessions (Table 1 – shaded) with varying waterlogging tolerance at the rosette stage were chosen to waterlog at the reproductive stage. Following 2 weeks of flowering, for each of the 8 accessions, six plants of the same accession were placed in 7.6-liter buckets (3 plants per bucket) and waterlogged for one week by filling the buckets with water up to the soil line at the top of the pots. An additional six plants of each accession were placed in buckets (3 plants per bucket), but watered normally, approximately every other day, so that the soil was completely saturated after watering, but dry before the next watering. The four buckets for each accession were arranged into two rows in a growth chamber, each containing one waterlogged and one control bucket and each of the two columns containing one waterlogged and one control bucket, so that buckets of the same treatment were not placed next to one another. After one week, the plants were taken out of the buckets and allowed to recover and were measured at maturity for total branch number, total plant height, reproductive plant height (total plant height minus height to first silicle), total silicle number, percentage of aborted silicles, average number of seeds per silicle for 10 silicles, total shoot dry weight, total seed weight, and days to maturity (or full senescence) after the start of flowering. For six of the accessions (950177, ISU_182, AMES_33895, ISU_002, ISU_077, and AMES_31012), an additional six plants were waterlogged and six plants were watered normally, and after 1 week they were immediately destructively harvested for biomass traits, including shoot and root fresh weight and total leaf number. Root and shoot tissues were then oven-dried at 50°C for 48 hours and root and shoot dry weights were also taken.

**Table 1.**
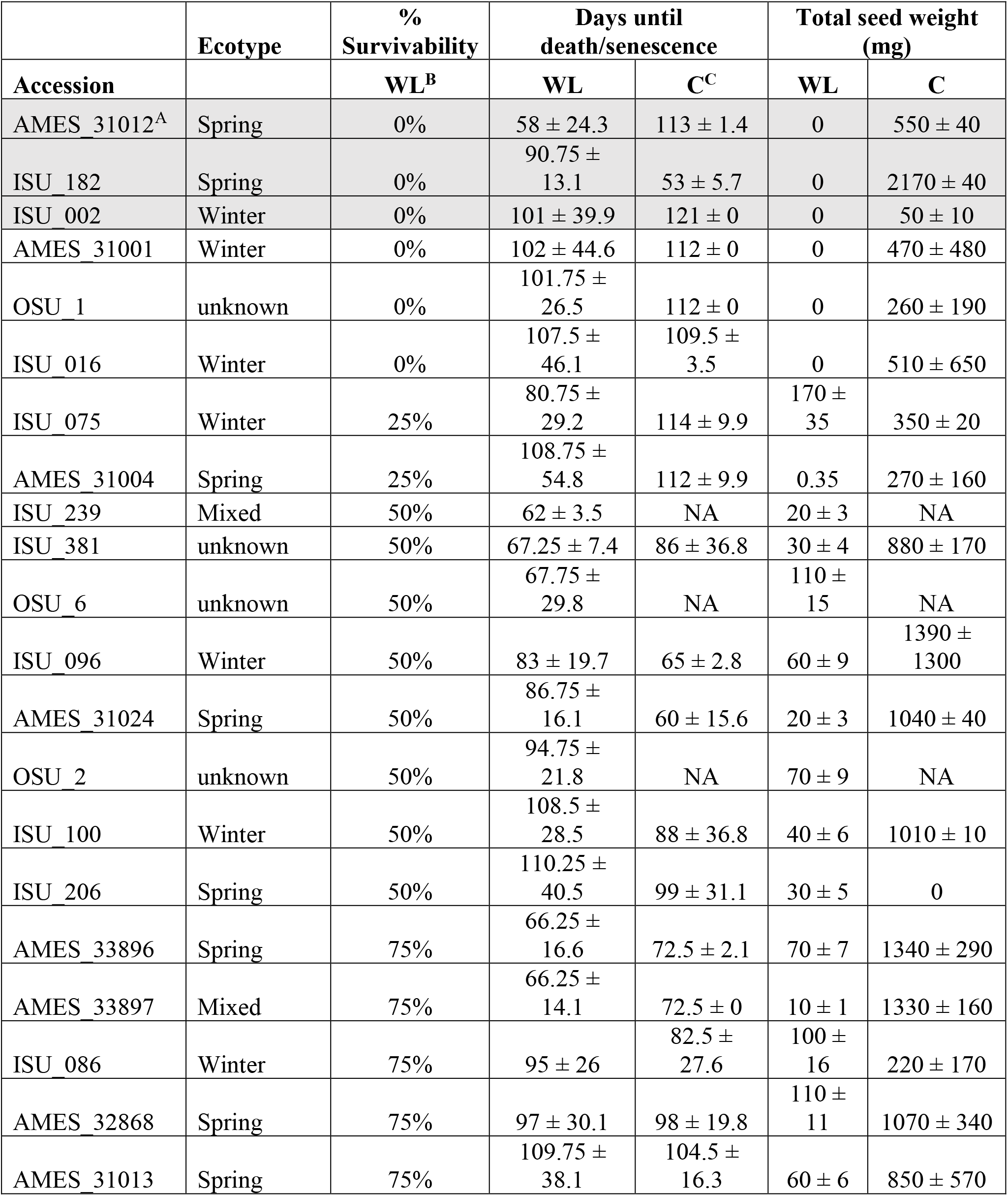
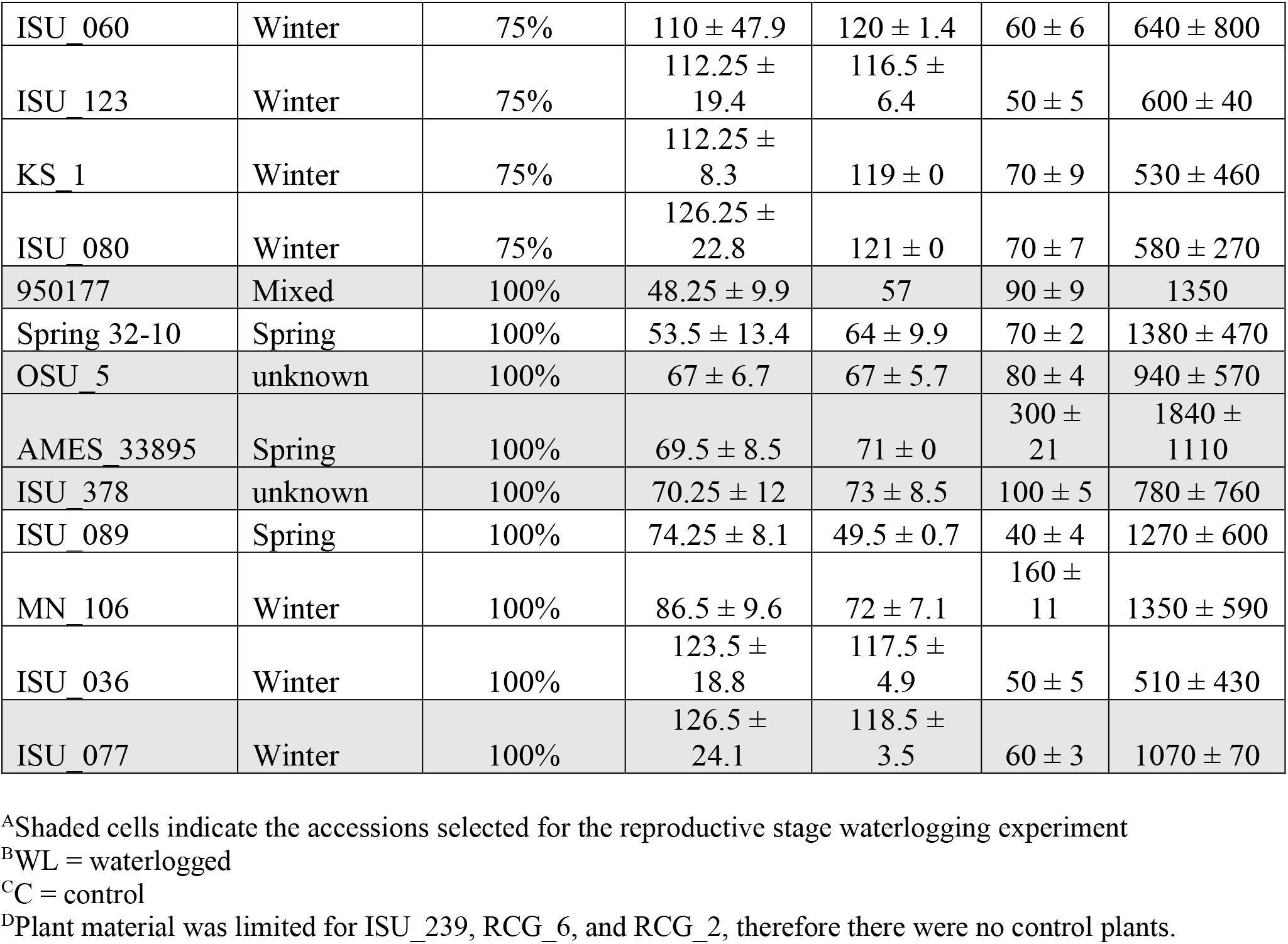
Percent survivability, number of days until plants died/senesced, and total seed weight under waterlogging at the rosette stage (mean ± standard deviation).

### Statistics

Normality and homogeneity of variance were tested with Shapiro-Wilks and Levene’s tests. Traits with unequal variances for at least one of the accessions were days to maturity, shoot dry weight, root dry weight, root fresh weight, seed weight, silicle number, and aborted silicle number. A Welch’s t-test was used to determine significance between waterlogged and control replicates of each measured trait for every accession waterlogged at the reproductive stage. A two-way analysis of variance (ANOVA) followed by a post-hoc Tukey’s Honestly Significant Difference (HSD) test with alpha = 0.05 was used to determine significance between waterlogged and control groups across accessions. A Pearson’s correlation coefficient was calculated to determine the relationship between SWA scores from region of origin and survivability, SWA score from region of origin and average seed weight, latitude from region of origin and survivability, and latitude from region of origin and average seed weight. All statistical analyses were conducted in RStudio (R version 4.2.1).

## Results

### Variation in survivability among natural accessions under prolonged rosette-stage waterlogging

To identify pennycress natural accessions that may be better adapted to wet and waterlogged environments, we assessed the climate data for the GPS coordinates of the 471 pennycress natural accession source populations to assign a spring soil water availability (SWA) score to each of the natural accessions (Figure 1A). About 37% of the accessions had low SWA scores (0-0.25), suggesting these accessions were collected from regions with drier spring climates, 57% had medium scores (0.25-0.65), and 6% had high scores (0.65-1), suggesting these accessions were collected from regions with wetter spring climates. Natural accessions with high SWA scores were collected from populations located in the Pacific Northwest U.S, Central Midwest U.S., Southeastern U.S., and Northern Europe, whereas natural accessions with low to moderate SWA scores were more widely distributed across the North American and European continents. We used the spring SWA scores to select a subset of pennycress natural accessions to experimentally test their performance under waterlogging stress. We selected 34 pennycress accessions with a range of low, medium, and high SWA scores from different geographic locations to capture accessions from regions with predicted differences in spring soil moisture content and to increase the likelihood of assessing genetically diverse populations (Supplemental Table 1 and Figure 1B). For instance, Armenian accessions were found to be significantly genetically diverse compared to accessions from the U.S., Europe, Canada, and the Caucasus region (Frels et al. 2019). We selected accessions from Canada, Sweden, Armenia, and across the U.S., including the Pacific Northwest, the Midwest, Southern, and Eastern states.

We waterlogged these 34 accessions at the rosette stage to determine survival rates. Out of the 34 accessions, six had 0% survivability, meaning all four replicates died, and nine accessions had 100% survivability, meaning all four replicates survived and produced seed under continuous waterlogging (Table 1). Out of the remaining 19 accessions, two had 25% survivability, eight had 50% survivability, and nine had 75% survivability. Plants from every accession showed signs of chlorosis, or leaf yellowing, which eventually led to tissue death in the plants that died (Supplemental Figure 1). Of the 25 accessions that had at least one of the four plant replicates die due to waterlogging stress, the plants of 24 of these accessions died while still in the rosette stage (Supplemental Figure 2A and B) and consisted of both winter and spring ecotype accessions. Only one accession, ISU_182, had all replicates die after flowering, but before seeds were produced (Supplemental Figure 2C). From the six accessions with 0% survivability, most died after an average of 100 days of continuous waterlogging, except for AMES_31012 which died after about 58 days (Table 1). This indicates that these pennycress accessions were relatively hardy to prolonged waterlogging at the rosette stage.

After 2 weeks of waterlogging, 47% of the accessions were bolting and 24% of those accessions were flowering, whereas the other half of the accessions were still in a rosette stage (Supplemental Figure 3). Accessions that were bolting and flowering after 2 weeks of waterlogging were comprised of 38% spring ecotypes and 13% winter ecotypes (remaining were unknown/mixed ecotypes), whereas those still in a rosette stage were 67% winter ecotypes and 28% spring ecotypes (1 accession was unknown). The waterlogged spring accessions with 100% survival flowered, reproduced, and matured quickly (on average 66 days), whereas the winter accessions with 100% survival flowered, reproduced, and matured more slowly (on average 112 days) (Figure 2A). Not surprisingly, since spring ecotypes flower quicker than winter ecotypes, overall, the spring accessions died or naturally matured on average around 84 days, whereas the winter accessions died or naturally matured on average around 105 days (Figure 2B). While the means for days to maturity/death were comparable between control and waterlogged plants for spring (82-84 days) and winter (105 days) ecotypes, the variation was higher in waterlogged compared to control plants by 7 days in the spring ecotypes and by 9 days in the winter ecotypes. The survivability of the accessions was not related to ecotype, since all levels of survivability (0%, 25%, 50%, 75%, and 100%) contained accessions from both winter and spring ecotypes (Table 1), however, some ecotypes were unknown or mixed (and excluded from above ecotype calculations).

**Figure 2.**
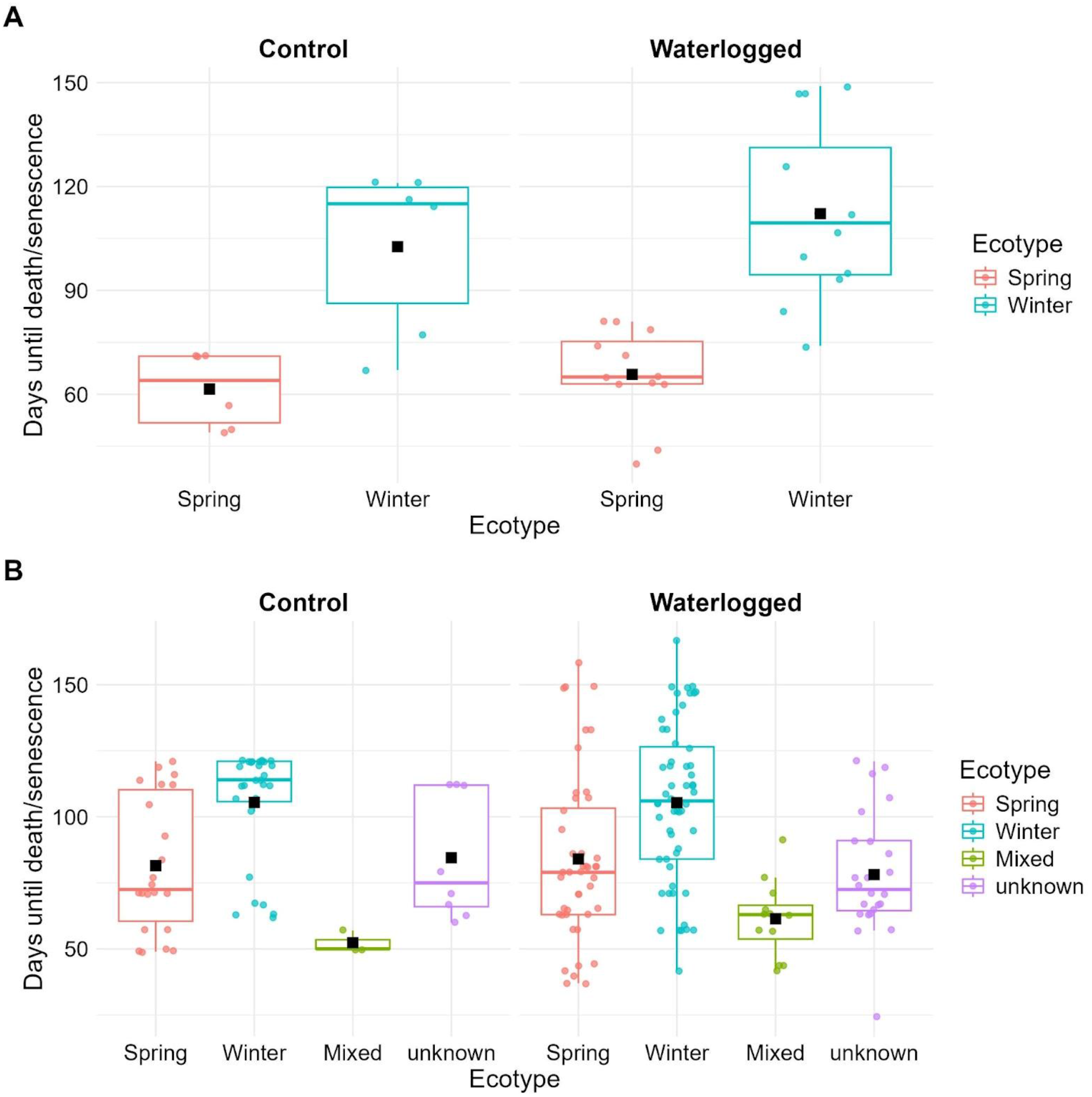
Boxplots showing maturity data for A) control and waterlogged rosette plants from only the accessions with 100% survival corresponding to spring and winter ecotypes and B) all control and waterlogged rosette plants from each accession corresponding to their ecotype. The black square represents the mean value.

Lastly, 28 of the accessions had at least one plant replicate that was able to produce seed under waterlogging, with the average seed weight ranging from 0.35-300mg (Table 1). The nine accessions with 100% survivability also had all four replicates produce seed, however, many of those accessions had reduced and aborted silicles compared to controls (Supplemental Figure 4), with an average seed weight of about 106mg for waterlogged accessions compared to 1,166mg for control accessions (Table 1).

### No significant correlations between geographic origin and plant fitness under rosette-stage waterlogging

It was hypothesized that survivability or average seed weight after waterlogging at the rosette stage would be correlated with the SWA scores or with latitudinal origin of the natural accessions. A weak positive correlation was revealed for SWA scores and average seed weight (R=0.36, Figure 3A). No correlation was discovered between SWA scores and survivability (R=0.18, Figure 3B), latitudinal origin and survivability (R=0.16, Figure 3C), and latitudinal origin and average seed weight (R=0.098, Figure 3D).

**Figure 3.**
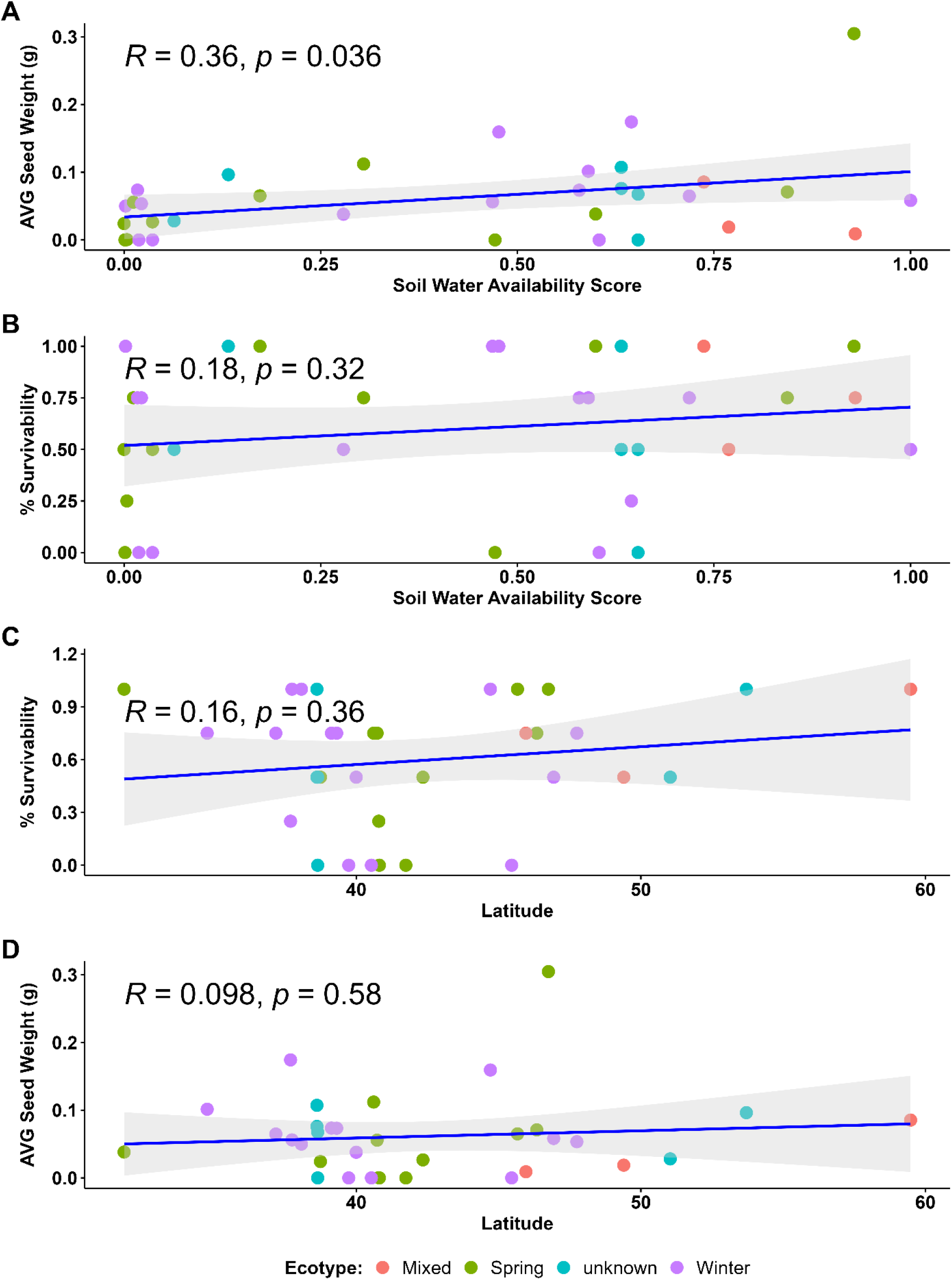
Pearson correlations between A) soil water availability score and percent survivability, B) soil water availability score and average seed weight, C) latitude and percent survivability, and D) latitude and average seed weight under waterlogging at the rosette stage. Accessions are color-coded by their ecotype.

### Variation in biomass among accessions immediately after reproductive stage waterlogging

Three waterlogging “susceptible” accessions and five waterlogging “tolerant” accessions were selected to be waterlogged for one week at the reproductive stage based on survivability, average seed weight, and the number of days it took for the plants to die/senesce under rosette-stage waterlogging (Table 1). Geographic location was also considered when selecting these eight accessions, to capture natural genetic variation between accessions. The susceptible (_S) accessions chosen were AMES_31012_S, ISU_002_S, and ISU_182_S. The tolerant (_T) accessions chosen were 950177_T, AMES_33895_T, ISU_077_T, ISU_378_T, and OSU_5_T.

Immediately following waterlogging, we destructively harvested plants to analyze various biomass traits among 6 of the accessions (two accessions were excluded from this analysis due to limited plant material) (Figure 4, Supplemental Table 2). Shoot fresh weight was significantly reduced by approximately 23-40% under waterlogging compared to the controls, and shoot dry weight was significantly reduced in AMES_33895_T by 23%, ISU_002_S by 37% and ISU_182_S by 25%. On the other hand, root fresh weight was significantly reduced in 950177_T (60%), AMES_33895_T (32%), ISU_077_T (40%), and AMES_31012_S (72%), but not in ISU_002_S and ISU_182_S. This could indicate a trade-off between shoot and root biomass in these latter two accessions. However, only AMES_31012_S had a significant 62% reduction in root dry weight under waterlogging. Furthermore, total leaf number was significantly reduced under waterlogging in ISU_077_T (42%), AMES_31012_S (31%), and ISU_182_S (26%). Immediately after waterlogging, there did not appear to be a relationship between biomass and rosette-stage tolerant or susceptible accessions, since most of the accessions had significant reductions in at least two or three biomass traits that varied between the accessions. For instance, shoot dry weight was reduced in the tolerant AMES_33895 accession, but also in the susceptible ISU_002 and ISU_182 accessions.

**Figure 4.**
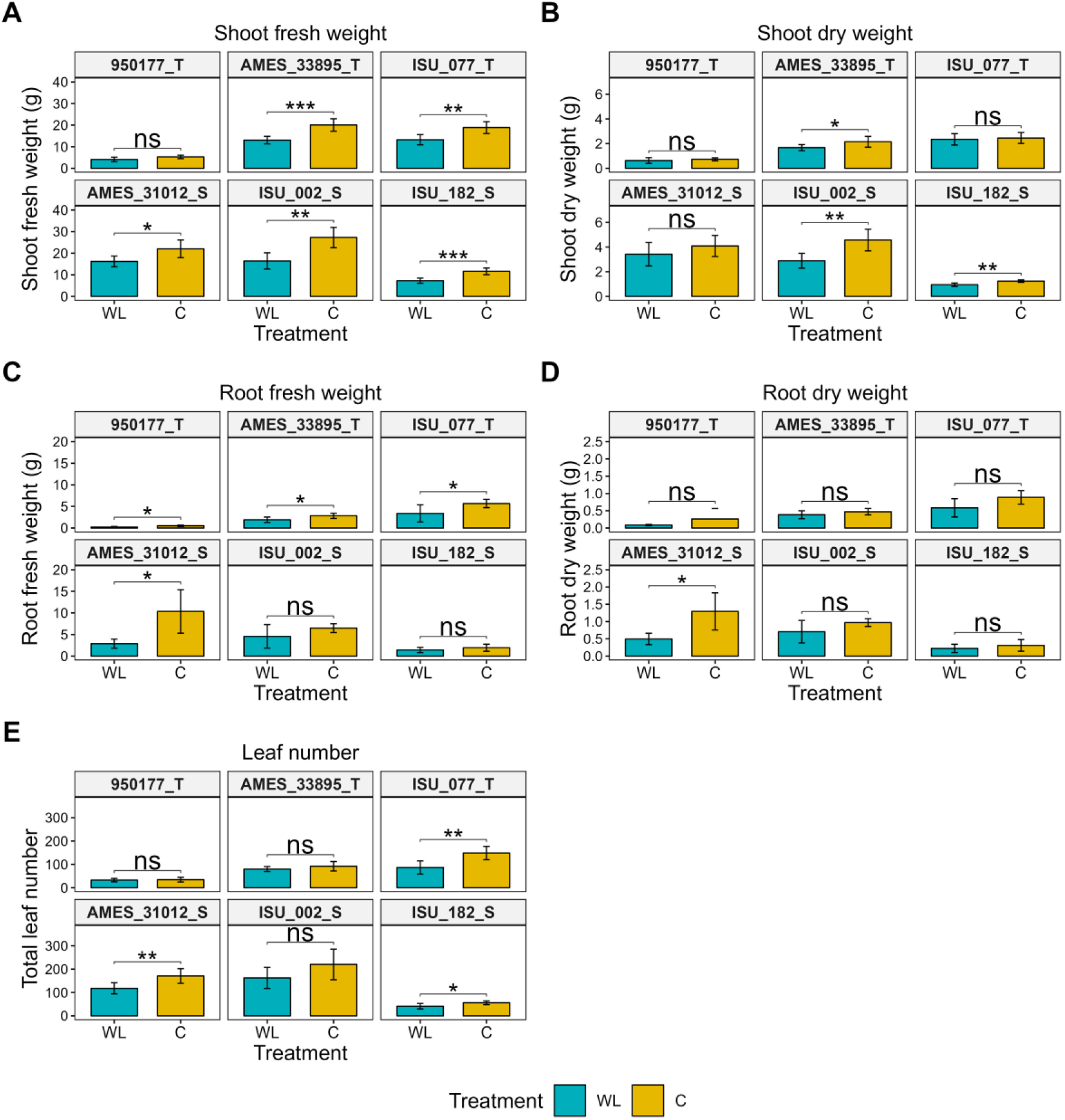
Biomass traits of 6 natural accessions immediately after waterlogging at the reproductive stage. A) Shoot fresh weight, B) Shoot dry weight, C) Root fresh weight, D) Root dry weight, and E) Total leaf number. C = Control and WL = Waterlogged. * = p-value < 0.05, ** = p-value < 0.01, ***=p-value < 0.001, ****=p-value < 0.0001.

There was also a significant interaction of treatment and accession for these biomass traits, as well as significant variation among the accessions (Supplemental Tables 2 and 3). For instance, the shoot fresh weight of 950177_T control plants (mean = 5.3g) was significantly lower compared to the other accessions (means = 11.6-27.3g). Root fresh and dry weight, as well as leaf number, of 950177_T control plants were significantly lower compared to ISU_077_T, AMES_31012_S, and ISU_002_S, as well. These results indicate that even under normal conditions 950177_T had low biomass overall compared to the other accessions tested, however, none of these biomass traits, except root fresh weight, were significantly reduced between waterlogged and control plants in 950177_T.

### Variation in morphology and seed yield among accessions at maturity following reproductive stage waterlogging

Once plants of the eight selected accessions recovered from one week of waterlogging at the reproductive stage and had fully matured/senesced, we collected several morphological traits (Figure 5, Supplemental Table 4). The total number of branches on waterlogged plants was significantly reduced by about half in AMES_33895_T (56%) and OSU_5_T (53%), and by about 75% in ISU_077_T and ISU_002_S. Plant height was significantly reduced in waterlogged plants for AMES_33895_T (8%), ISU_077_T (25%), ISU_378_T (12%), AMES_31012_S (17%), and ISU_182_S (23%). Reproductive plant height was also significantly reduced in waterlogged plants by 20-41% in AMES_33895_T, ISU_077_T, ISU_378_T, AMES_31012_S, and ISU_002_S. Total shoot dry weight was significantly reduced in six of the eight waterlogged accessions, except for 950177_T and AMES_31012_S, compared to the controls. Lastly, we looked at the number of days it took for the plants to mature after the initial start of flowering, revealing 950177_T, ISU_077_T, ISU_378_T, AMES_31012_S, and ISU_002_S all matured significantly more quickly than the controls, with some maturing on average about 4-9 days quicker than controls (950177_T, ISU_378_T, and AMES_31012_S) and some maturing on average about 17-24 days quicker (ISU_077_T and ISU_002_S).

**Figure 5.**
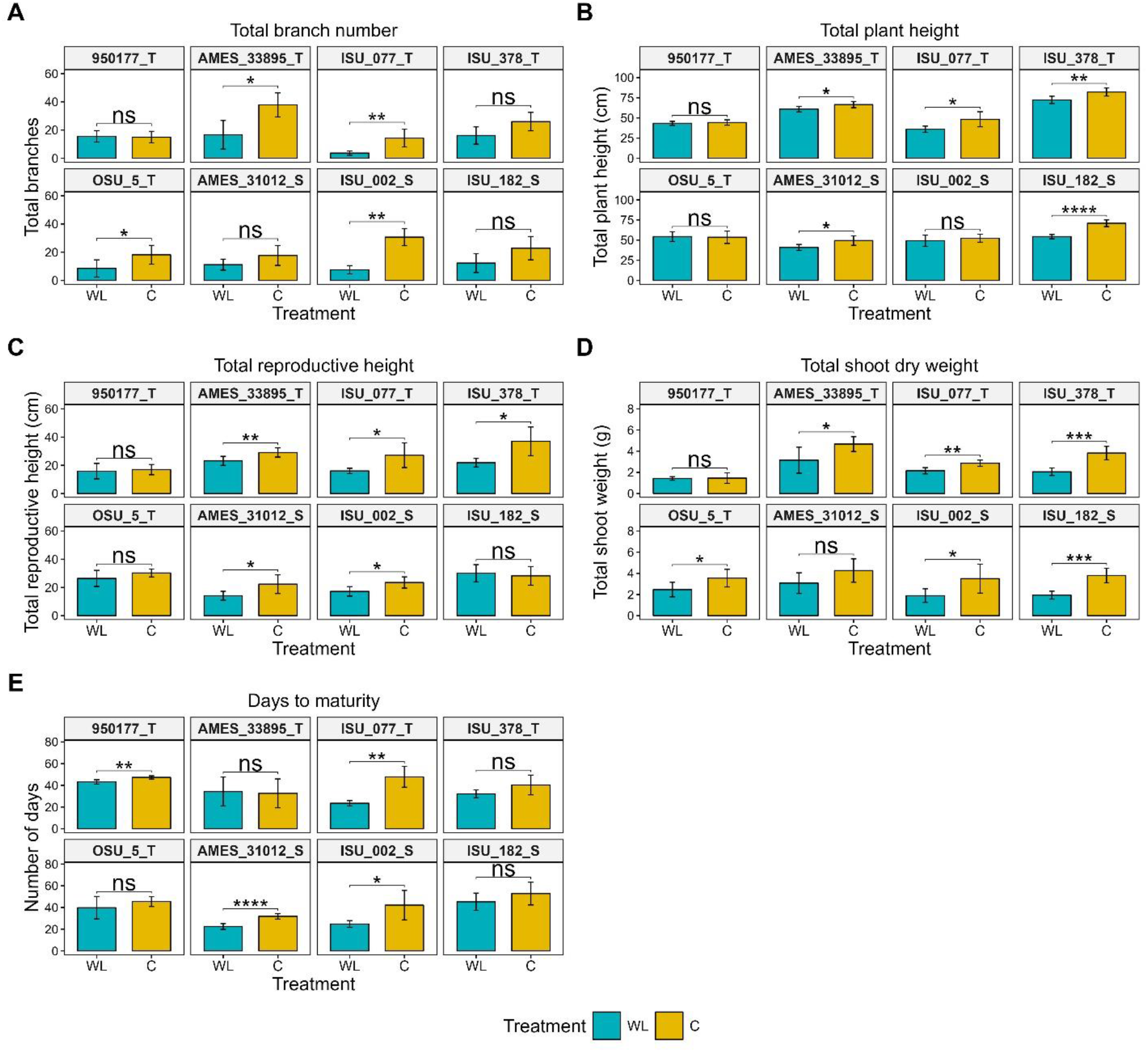
Morphological traits of 8 natural accessions collected at the time of maturity, after recovery from 1 week of waterlogging at the reproductive stage. A) Total branch number, B) Total plant height, C) Total reproductive height, D) Total shoot dry weight, E) Days to maturity. C = Control and WL = Waterlogged. * = p-value < 0.05, ** = p-value < 0.01, ***=p-value < 0.001, ****=p-value < 0.0001.

Different components of seed yield were also measured, such as silicle number, the percentage of aborted silicles, the average number of seeds per silicle, and total seed weight (Figures 6 and 7, Supplemental Table 5). The total number of silicles was significantly reduced by over half compared to controls (53-77%) in six accessions under waterlogging, but not in 950177_T and OSU_5_T. The number of aborted silicles, or silicles that did not fully develop to contain seeds, was significantly increased under waterlogging by approximately 43% for 950177_T, AMES_33895_T, and ISU_182_S. The number of seeds per silicle stayed about the same under waterlogging compared to controls in most accessions, except for ISU_077_T, which showed a 19% reduction. Seed weight was significantly reduced in waterlogged plants in 6 of the accessions: AMES_33895_T (50%), ISU_077_T (71%), ISU_378_T (45%), AMES_31012_S (71%), ISU_002_S (82%), and ISU_182_S (57%). The accessions without a significant impact on total seed weight under waterlogging were 950177_T and OSU_5_T, where waterlogged plants had reduced seed yield by only 14% and 20%, respectively. Reductions in yield and morphological traits might be tied to quicker maturity observed in several of the accessions. For instance, ISU_378_T and AMES_31012_S, which matured 4-9 days quicker than controls, had significant reductions in plant height, silicles, and total seed weight, and ISU_077_T and ISU_002_S, which matured 17-24 days quicker, also showed a significant reduction in these traits, as well as in branch number and shoot dry weight. Early maturity does not appear to be the only factor effecting yield traits, though, since AMES_33895_T and ISU_182_S did not have significant differences in days to maturity between waterlogged and control plants, but still had a significant reduction in silicle number and total seed weight, and an increase in aborted silicles. Furthermore, based on morphology and yield data, both susceptible and tolerant accessions had multiple traits negatively impacted by waterlogging, though yield (silicle number and seed weight) was significantly negatively impacted by waterlogging in all three susceptible accessions, but was not significantly affected in two of the tolerant accessions (950177_T and OSU_5_T). These results indicate that, even though the “tolerant” lines were able to survive prolonged waterlogging at the rosette stage of development, most of these accessions were sensitive to waterlogging at the reproductive stage, in that it resulted in reduced yield and biomass. Additionally, there were not any traits that were specifically negatively impacted by waterlogging only in the tolerant or susceptible accessions.

**Figure 6.**
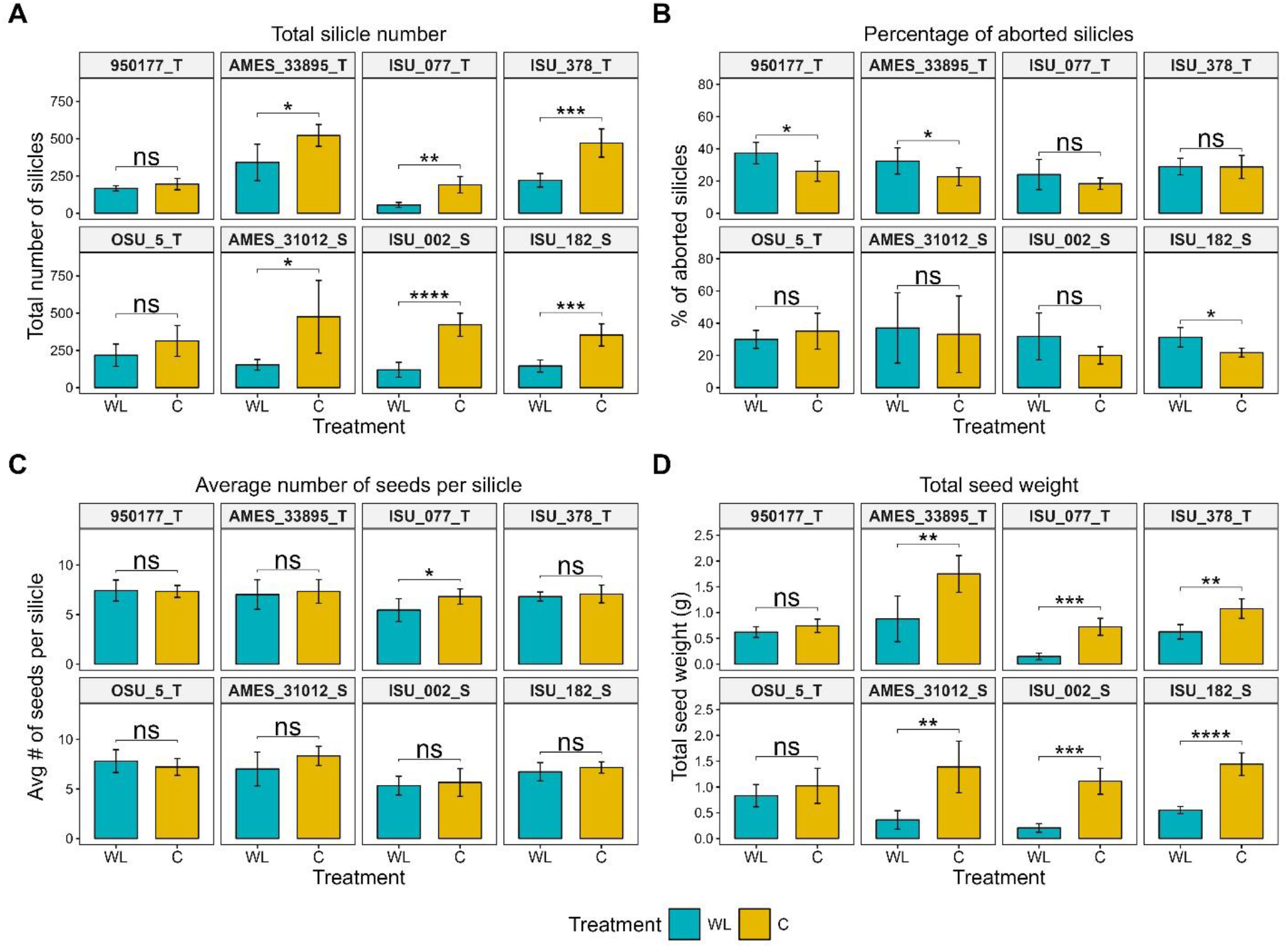
Yield traits of 8 natural accessions collected at time of maturity, after recovery from 1 week of waterlogging at the reproductive stage. A) Total silicle number, B) Percentage of aborted silicles, C) Average number of seeds per silicle, D) Total seed weight. C = Control and WL = Waterlogged. * = p-value < 0.05, ** = p-value < 0.01, ***=p-value < 0.001, ****=p-value < 0.0001.

**Figure 7.**
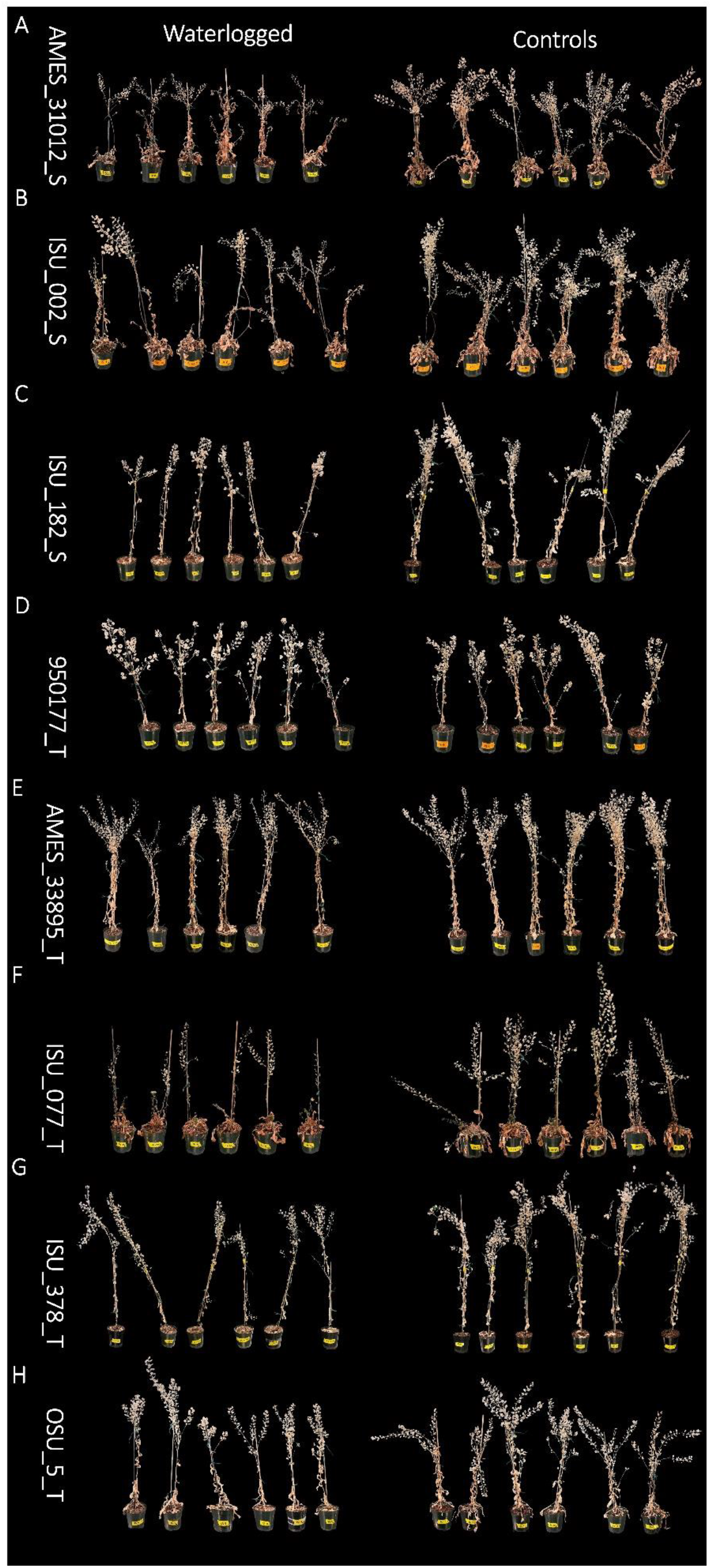
Pennycress waterlogged (1 week at the reproductive stage) and control replicates at time of maturity. A) AMES_31012_S, B) ISU_002_S, C) ISU_182_S, D) 950177_T, E) AMES_33895_T, F) ISU_077_T, G) ISU_378_T and H) OSU_5_T.

We also examined variation in morphological and yield traits between accessions (Supplemental Tables 6 and 7). There was a significant interaction between treatment and accession for all morphological and yield traits except for shoot dry weight, the percentage of aborted silicles, and the number of seeds per pod, indicating that the impact of waterlogging on most of these traits differs significantly across accessions. Additionally, there was significant variation among the accessions for all of these traits except for the percentage of aborted silicles. For instance, ISU_378_T control plants were significantly taller than all the other accessions with an average height of 82cm and 950177_T control plants had significantly lower shoot dry weight than all the other accessions, except for ISU_077, with an average weight of 1.5g. Therefore, in addition to natural phenotypic variation among pennycress accessions in response to waterlogging, there is also natural variation among accessions under normal conditions.

*The 950177 accession was tolerant to waterlogging, but had naturally low biomass and yield* Waterlogging at the reproductive stage revealed that 950177 performed well under waterlogging stress without a significant impact on seed yield. However, we found that this accession produced significantly lower biomass and seed yield compared to the other accessions. For instance, control plants had an average shoot weight of 1.5g at the time of maturity, whereas all other accessions had shoot weights greater than 2.9g (Supplemental Table 4). Additionally, average total seed weight in the 950177 controls was 0.7g, whereas the other accessions, except ISU_077, had average total seed weights over 1g (Supplemental Table 5). These differences in biomass can be visually observed in Figure 7. Therefore, while 950177 is tolerant to waterlogging, it has low seed yield.

## Discussion

### Variation in survival under rosette stage waterlogging

Our results showed variation in the survivability of pennycress under prolonged waterlogging imposed at the rosette stage, where half of the accessions had 75-100% survival. For winter ecotype accessions, this developmental stage corresponds with the winter season (late November – late March) where the plants are dormant until temperatures rise in the spring. In winter ecotype plants, prolonged exposure to cold temperatures slows plant metabolism, a process called vernalization, but accelerates flowering time when environmental conditions are favorable for reproduction (Dorn et al. 2018; Kim et al. 2009). We predicted that winter accessions might respond to waterlogging similar to cold stress by remaining in a quiescent state until conditions were favorable again, whereas spring accessions might quickly flower and reproduce under waterlogging using an escape strategy (Akman et al. 2012; Kazan and Lyons 2016; Müller et al. 2021). However, the average days to maturity/death were comparable between control and waterlogged plants in the spring (82-84 days) and winter ecotypes (105 days). While this would indicate there is no unique survival mechanism under waterlogging based on ecotype, we did observe increased variation in days to maturity of both ecotypes under waterlogging stress, suggesting different survival strategies may have evolved in some populations despite ecotype. Roughly half of all winter and spring accessions had more than or equal to 75% survival, indicating that some accessions from both winter and spring ecotypes had similar survival success. Additionally, factoring in early maturing vs late maturing accessions under waterlogging (based on a total average of 90 days), both groups had a total of 9 accessions that had 3-4 reps flower and produce seed, indicating both early and late maturing accessions could survive and reproduce under waterlogged conditions, so days to maturity was not indicative of survival. Further investigation of early maturing and late maturing accessions did show differences in survival strategy between ecotypes. Of the six early maturing spring accessions, 66% had 75-100% survivability under waterlogging, whereas only one of the three early maturing winter lines had 75-100% survivability (33%). For the late maturing accessions, two of five (40%) spring accession had 75-100% survivability under waterlogging, whereas seven out of 11 (63%) of the winter lines had 75-100% survivability. These data suggest, though spring and winter ecotypes can be both early and late maturing under waterlogging, a greater proportion of spring ecotypes that matured early had higher survivability, and a greater proportion of winter ecotypes that matured later had higher survivability. Not surprisingly there was a tradeoff in days to maturity and seed weight. On average, the spring and winter accessions with 75-100% survivability had higher total seed weight in the early maturing group (128mg) than the late maturing group (69mg). This is likely due to the catabolism of primary metabolites for energy by late maturing plants for prolonged survival under waterlogged conditions, whereas early maturing plants likely allocated those resources to the seeds for reproduction. In our rosette waterlogging study we purposefully waterlogged plants until they died or senesced, but we hypothesize that if the plants were allowed to recover after a couple weeks of waterlogging at the rosette stage, the results would be different, and the accessions demonstrating the quiescent strategy, delaying maturity by remaining dormant as rosettes under waterlogging, would have higher seed yield than the early escape strategy accessions at the time of maturity by replenishing carbohydrates via photosynthesis once the water recedes to allocate to reproductive tissues. Testing the survival and yield of plants after a recovery period following waterlogging at the rosette stage would test these hypotheses.

The variation in pennycress survival under rosette-stage waterlogging hints at adaptation to local environments across these accessions. Variation in waterlogging and submergence responses has been reported for natural accessions of Arabidopsis (Pigliucci and Kolodynska 2002; Vashisht et al. 2011). For example, 86 natural Arabidopsis accessions had significant variation in submergence tolerance, with northern accessions correlated with better survivability, which could indicate they were better adapted to wetter environments (Vashisht et al. 2011). Since pennycress typically flowers in the spring and likely encounters fluctuations in spring flooding based on geography, each pennycress population/accession likely evolved to adapt to the type of flooding common at the source population location. Therefore, we hypothesized accessions with lower SWA scores would have lower survivability under prolonged waterlogging as rosettes and accessions with higher SWA scores would have higher survivability, due to being better adapted to wetter environments. We tested if the SWA scores were correlated with waterlogging survival or yield after prolonged waterlogging at the rosette stage, but only observed a weak positive correlation between SWA scores and average seed weight. We don’t know how long these pennycress populations have been established in their source environments, so one reason for the weak positive correlation could be that some of these populations were newly established as a result of recent dispersal events. Testing more accessions might provide better resolution into whether geographic location and predicted water availability are correlated with flooding tolerance phenotypes. It is also important to note the limitations of the climate data used in this study. For instance, only the closest coordinate for which climate data was available was used, and although the latitudes or longitudes of an accession source population were no more than 0.1° away from the climate coordinate, there could still be natural topographic differences, such as hills or ditches, that were not taken into account. We also investigated the survivability of waterlogged accessions with the highest (0.7-1) SWA scores. 71% of the accessions with SWA scores between 0.7-1 had 75% or 100% survivability. As a reference, only 36% of accessions with SWA scores between 0-0.3 had 75% or 100% survivability. These results support that accessions collected from predicted wetter environments had greater survivability under prolonged waterlogging. To get a better idea of the relationship between soil moisture of the geographic origins of these pennycress natural accessions and their waterlogging tolerance, precise soil moisture readings could be obtained using sensors in the exact coordinates where these accessions were collected. This would provide further support that variation in pennycress waterlogging tolerance is attributed to local adaptation to environments with different water availability.

Pennycress survived for multiple weeks under prolonged rosette-stage waterlogging, where the first accessions to die or mature lived for 7-8 weeks under waterlogged conditions. Because these pennycress accessions lasted multiple weeks under rosette waterlogging, and many accessions were able to produce seed while being continuously waterlogged, we can conclude that many of these pennycress accessions are relatively fit in waterlogged environments. However, among the accessions that survived, seed yield was reduced compared to the controls. Other *Brassicaceae* crops under prolonged waterlogging during seedling and vegetative development also had negative effects on yield and biomass. For instance, *B. napus* (cv. 601) exposed to prolonged waterlogging (30d) in a field environment at the seedling and floral bud appearance stages had a significant reduction in seed yield (Zhou and Lin 1995). Additionally, *B. oleraceae* (var. *capitata* and *italica*) exposed to prolonged waterlogging (25-45d) beginning at the seedling stage revealed reduced biomass and inhibited flower production (Casierra-Posada and Cutle 2017; Casierra-Posada and Peña-Olmos 2022). Currently, there are very few reports of how other *Brassicaceae* species perform when grown under prolonged waterlogging at early stages of development, including the rosette stage, therefore it’s not clear how the variation in morphological and yield traits observed in these pennycress populations relate to other *Brassicaceae* relatives.

### Biomass was negatively impacted in several accessions immediately after waterlogging at the reproductive stage

The rosette stage waterlogging experiment determined pennycress survivability under prolonged waterlogging to inform the selection of accessions to further study at the reproductive stage, which occurs in the spring when pennycress fields are most susceptible to waterlogged soil following heavy rainfall. By testing three tolerant and three susceptible pennycress accessions under reproductive stage waterlogging for 1 week, we found variation in biomass traits immediately after waterlogging, both between controls and waterlogged plants and between accessions. Unlike many of the other accessions, 950177_T did not have any significant reductions in shoot dry weight, root dry weight, or leaf number. Many of the accessions did not have significant changes in root dry weight except for AMES_31012_S. Similarly, no significant changes in root dry weight immediately after waterlogging were previously observed in MN106 and SP32-10 (Combs-Giroir et al. 2024), however, we expected that some accessions would have reductions in root dry weight following waterlogging, since reduced root biomass is a common observation under flooding in other closely related species. For instance, reduced root and shoot weight was reported in the closely related oilseed crop, *B. napus*, following waterlogging imposed during reproductive stages (Liu et al. 2020; Liu and Zwiazek 2022; Ploschuk et al. 2023; Ploschuk et al. 2021; Ploschuk et al. 2020; Ploschuk et al. 2018; Wollmer, Pitann and K. H. Mühling 2018). A significant reduction in root dry weight was observed at maturity for six out of the eight pennycress accessions tested, though, suggesting it may take time following waterlogging to reallocate assimilates stored in the roots. Root biomass is commonly reduced following flooding stress as a result of carbohydrates either being consumed during fermentation under hypoxia, or as a result of expending carbohydrates for shoot growth to escape floodwaters, as is the case with some species when under submergence (Akman et al. 2012; Combs-Giroir and Gschwend 2024; Müller et al. 2021; Sasidharan et al. 2013). Lastly, the susceptible accessions did not consistently have significant reductions in all biomass traits, and some tolerant accessions did have significant reductions in biomass traits, indicating that rosette-stage waterlogging tolerance does not necessarily mean that biomass would not be impacted immediately after reproductive-stage waterlogging. Therefore, we also looked at yield traits at maturity to see if rosette-stage tolerance aligned with reproductive-stage tolerance in terms of yield.

### Six accessions had reduced yield following reproductive stage waterlogging

After plants had recovered from 7 days of waterlogging at the reproductive stage and naturally senesced, several morphology and yield traits were collected. Total seed weight was significantly reduced in every accession, except 950177_T and OSU_5_T. Lower total seed weight could be attributed to significantly lower silicle numbers in the six accessions, as well as lower branch numbers in half of these accessions. Lower seed weight, silicle numbers, and branching likely significantly contributed to the reduced shoot dry weight in five of the six accessions (except AMES_31012_S). This is not a surprising result, since plants typically prioritize using resources for stress and defense responses at the expense of growth (Zhang et al. 2020). This response was also observed for *B. napus*, a closely related oilseed crop, which experienced reduced seed yield following reproductive-stage waterlogging (Ploschuk et al. 2023; Ploschuk et al. 2021; Ploschuk et al. 2018; Wollmer, Pitann and K. H. Mühling 2018; Xu et al. 2015). Since many of the pennycress accessions were negatively impacted by reproductive-stage waterlogging as it related to morphology and yield, we can conclude that although some accessions can survive and even reproduce following prolonged waterlogging, it does not mean that seed yield of those accessions won’t be significantly impacted. In this study, tolerance at the rosette stage was defined as accessions that were able to survive and produce seed under prolonged waterlogging, though it should be noted that all of the 34 accessions showed reduced seed yield under rosette waterlogging compared to controls. Tolerance at the reproductive stage was defined as accessions with minimal impacts on morphology and seed yield at maturity after one week of waterlogging. Energy expenditure and resource allocation during the vegetative versus reproductive stage are already different under optimal growing conditions, but additional hypoxic stress can further affect resource use and allocation, impacting growth and development at these stages. Waterlogged tissues have limited access to oxygen, inhibiting oxidative phosphorylation and significantly decreasing ATP production. To continue to support ATP production until the water recedes, resources are invested in fermentation and glycolysis to produce energy for basic biological functions and survival rather than being allocated to seed production (Combs-Giroir et al. 2024). However, reproductive-stage waterlogging did not have a significant impact on the seed yield of two accessions, suggesting these accessions may have evolved mechanisms to balance survival and reproduction under waterlogging stress.

### Two accessions were tolerant to reproductive stage waterlogging

Unlike many of the other accessions, 950177_T did not have any significant reduction in shoot dry weight, root dry weight, or leaf number immediately after waterlogging, and along with OSU_5_T, it did not have significant reductions in morphology or yield traits at maturity. However, under normal growth conditions, this accession had significantly lower shoot and root biomass and seed yield compared to many of the other accessions tested in this study and those previously tested (MN106 and SP32-10) (Combs-Giroir et al. 2024). For instance, the average seed weight of MN106 and SP32- 10 plants ranged from 1.2-1.4g, whereas the average seed weight of 950177_T was only 0.7g. This might suggest that this accession minimizes resource allocation for vegetative and reproductive growth to provide more resources to respond to abiotic and biotic stress, such as storing carbohydrates or producing secondary metabolites (Bechtold et al. 2018; Figueroa-Macías et al. 2021). Interestingly, this accession, originating in Stockholm, Sweden, had the highest latitude out of all 34 original accessions tested and had a high SWA score (0.737). Therefore, this would be an accession worth testing for tolerance to other abiotic stresses, such as cold or freezing stress, and seeing if any physical and molecular responses overlap with waterlogging stress.

When breeding for waterlogging tolerance, the maintenance of seed yield is the most important factor to consider, as well as ensuring there are no significant impacts on seed or oil quality. While 950177_T had no impacts on seed yield after waterlogging, it might not be an ideal parental line to breed for polygenic traits like yield and waterlogging tolerance since yield is much lower than those used in breeding programs, like MN106. Furthermore, analysis of oil content and fatty acid constituents should be evaluated in 950177_T and OSU_5_T to see if there are any negative impacts on oil yield and quality following waterlogging events. Another trait to consider when breeding for waterlogging tolerance is plant height. As a side effect of flooding, plants are more prone to lodging, therefore, accessions with tall plant heights, such as ISU_182 and ISU_378, would not be desirable from an agronomic standpoint. Another approach for integrating waterlogging tolerance into elite, high-yielding varieties is by identifying genes contributing to waterlogging tolerance phenotypes and using a gene editing approach to maintain the genetic background of high-yielding lines but incorporating alleles contributing to waterlogging tolerance. The 950177_T and OSU_5_T accessions, which were tolerant to both rosette and reproductive stage waterlogging, can be used for further investigation of genetic variation contributing to waterlogging tolerance. This can be accomplished via comparisons with waterlogging susceptible accessions, such as ISU_002_S and ISU_182_S, for QTL analysis or various comparative transcriptomic, metabolomic, or proteomic analyses. This will help identify significant genetic differences between tolerant and susceptible accessions and can be followed by functional validation of candidate genes and integration into commercial varieties.

## Conclusions

Based on our results, there is natural variation in waterlogging tolerance in natural pennycress accessions at the rosette and reproductive stages, with a weak positive correlation between the spring water availability at their respective geographic origins and seed weight. We have identified two accessions tolerant to reproductive stage waterlogging, which can be used in comparison with susceptible lines identified here to further investigate the genes and molecular mechanisms behind pennycress waterlogging tolerance and inform breeding waterlogging-resilient pennycress. Furthermore, screening the whole pennycress germplasm curated by IPReP for phenotypic responses to waterlogging, such as leaf chlorosis/senescence, leaf area index, or seed yield, could also aid the identification of other waterlogging tolerant natural accessions and contribute to the identification of candidate genes through genome-wide association studies. A potential way to test this is through a large-scale greenhouse or field study with coordinated high-throughput phenotyping methods, such as drone imaging and leaf scans. This approach would help identify traits, such as accelerated senescence, that might correlate with ecotypes or waterlogging resilience.

## Data Availability

The climate data used for this study can be accessed via the TerraClimate repository (https://www.climatologylab.org/terraclimate.html). Custom scripts for the climate analyses can be found on Github (https://github.com/jlager/pennycress-soil-moisture). The data collected from the experiments of this study are summarized in the main and supplemental tables and figures of this manuscript.

## Author Contributions

RCG and ARG conceptualized the project. JL and DAJ performed climate analysis. RCG performed waterlogging experiments, data collection, data analyses, and data visualization. RCG and ARG interpreted the data. RCG wrote the initial manuscript draft. RCG, JL, DAJ, and ARG edited the manuscript. ARG supervised and provided funding for the project.

## Funding

This research was funded by the U.S. Department of Energy, Office of Science, Office of Biological and Environmental Research, Genomic Science Program [DE-SC0021286]. Combs-Giroir was also financially supported in part by the National Science Foundation Graduate Research Fellowship Program.

## Acknowledgements

We would like to thank the numerous citizen scientists who assisted with collecting seeds for the IPReP collection and acknowledge Maliheh Esfahanian, Barsanti Gautam, Donald Wyse, Winthrop Phippen, Ratan Chopra, and John Sedbrook for organizing the collection. We also acknowledge the IPReP team for collectively bulking seed from the natural accessions and for feedback and direction related to this project. We would like to acknowledge Gary Posey, Thiranya Wanigarathna, Alex Solum, Molly Dougherty, Rosemary Ball, and Tara Creech for assistance with plant care & maintenance and/or data collection.

## Notes

### Competing Interest Statement

The authors have declared no competing interest.

