## Supplemental Tables and Figures for "Natural variation in physical responses to waterlogging across climate-diverse pennycress accessions"

**Supplemental Table 1.** List of accessions used for screening rosette-stage plants for survivability under waterlogging.

| Accession | Location | Soil Water Availability Score | Ecotype | Available |
|---|---|---|---|---|
| AMES_31024 | Nathrop, Colorado, USA | 0.000 | Spring | GRIN |
| AMES_31012[A] | Grover, Colorado, USA | 0.001 | Spring | GRIN |
| ISU_036 | Beulah, Colorado, USA | 0.002 | Winter | ISU - Sedbrook |
| AMES_31004 | Nunn, Colorado, USA | 0.004 | Spring | GRIN |
| AMES_31013 | La Porte, Colorado, USA | 0.012 | Spring | GRIN |
| KS_1 | Oakley, Kansas, USA | 0.017 | Winter | ISU - Sedbrook |
| AMES_31001 | Ft. Collins, Colorado, USA | 0.019 | Winter | GRIN |
| ISU_123 | Malta, Missouri, USA | 0.022 | Winter | ISU - Sedbrook |
| ISU_206 | Hemingford, Nebraska, USA | 0.036 | Spring | ISU - Sedbrook |
| ISU_002 | Harding County, South Dakota, USA | 0.036 | Winter | ISU - Sedbrook |
| ISU_381 | Southeast Calgary, Calgary, Alberta, CAN | 0.064 | unknown | ISU - Sedbrook |
| ISU_378 | Sturgeon County, Albertab, CAN | 0.133 | unknown | ISU - Sedbrook |
| Spring 32-10 | Bozeman, Montana, USA | 0.173 | Spring | ABRC |
| ISU_100 | Saint Joseph, Pennsylvania, USA | 0.279 | Winter | ISU - Sedbrook |
| AMES_32868 | Aragatsotn Province, Armenia | 0.305 | Spring | GRIN |
| ISU_077 | Highland, Virginia, USA | 0.469 | Winter | ISU - Sedbrook |
| ISU_182 | North Windham, Connecticut, USA | 0.472 | Spring | ISU - Sedbrook |
| MN106 | Coates, Minnesota, USA | 0.476 | Winter | ABRC |
| ISU_080 | Athens, Ohio, USA | 0.579 | Winter | ISU - Sedbrook |
| ISU_086 | Colbert, Alabama, USA | 0.590 | Winter | ISU - Sedbrook |
| ISU_089 | Troy, Alabama, USA | 0.599 | Spring | ISU - Sedbrook |

| | | | | |
|---|---|---|---|---|
| ISU_016 | Jacksonville, Illinois, USA | 0.604 | Winter | ISU - Sedbrook |
| OSU_5 | Marthasville, Missouri, USA | 0.632 | unknown | OSU - Collected in this study |
| OSU_6 | Marthasville, Missouri, USA | 0.632 | unknown | OSU - Collected in this study |
| ISU_075 | Myrphy Sboro, Illinois, USA | 0.645 | Winter | ISU - Sedbrook |
| OSU_1 | Marthasville, Missouri, USA | 0.653 | unknown | OSU - Collected in this study |
| OSU_2 | Marthasville, Missouri, USA | 0.653 | unknown | OSU - Collected in this study |
| ISU_060 | Green County, Missouri, USA | 0.719 | Winter | ISU - Sedbrook |
| 950177 | Sollentuna, Sweden | 0.737 | Mixed | ABRC |
| ISU_239 | Hope, British Columbia, CAN | 0.769 | Mixed | ISU - Sedbrook |
| AMES_33896 | Culdesac, Idaho, USA | 0.843 | Spring | GRIN |
| AMES_33895 | Moscow, Idaho, USA | 0.928 | Spring | GRIN |
| AMES_33897 | Grangeville, Idaho, USA | 0.930 | Mixed | GRIN |
| ISU_096 | Onaway, Idaho, USA | 1.000 | Winter | ISU - Sedbrook |

[A]Shaded rows indicate the accessions chosen for reproductive stage waterlogging experiment

**Supplemental Table 2.** Means and standard deviations of biomass traits immediately after waterlogging at the reproductive stage in six natural accessions.

| Accession | | Shoot FW[C] (g) | p-val[D] | Shoot DW[E] (g) | p-val | Root FW (g) | p-val | Root DW (g) | p-val | Leaf Number | p-val |
|---|---|---|---|---|---|---|---|---|---|---|---|
| **950177_T[A]** | C[B] | 5.3 ± 0.8$^{f}$ | 0.06 | 0.7 ± 0.1$^{g}$ | 0.38 | 0.5 ± 0.2$^{d}$ | 0.01* | 0.3 ± 0.3$^{def}$ | 0.22 | 34.3 ± 9.9$^{ef}$ | 0.75 |
| | WL | 4.1 ± 1.1$^{f}$ | | 0.6 ± 0.2$^{g}$ | | 0.2 ± 0.1$^{d}$ | | 0.1 ± 0.0$^{f}$ | | 32.7 ± 7.4$^{f}$ | |
| **AMES_33895_T** | C | 20.1 ± 2.9$^{bc}$ | 0* | 2.2 ± 0.4$^{def}$ | 0.05* | 2.8 ± 0.6$^{bcd}$ | 0.03* | 0.5 ± 0.1$^{cdef}$ | 0.18 | 91.8 ± 20.3$^{cde}$ | 0.24 |
| | WL | 13.1 ± 1.7$^{d}$ | | 1.7 ± 0.2$^{efg}$ | | 1.9 ± 0.6$^{cd}$ | | 0.4 ± 0.1$^{def}$ | | 79.8 ± 10.6$^{def}$ | |
| **ISU_077_T** | C | 18.9 ± 2.7$^{bc}$ | 0* | 2.5 ± 0.4$^{cde}$ | 0.7 | 5.7 ± 0.9$^{b}$ | 0.04* | 0.9 ± 0.2$^{abc}$ | 0.05 | 148.5 ± 28.5$^{bc}$ | 0* |
| | WL | 13.2 ± 2.4$^{d}$ | | 2.4 ± 0.5$^{de}$ | | 3.4 ± 2$^{bcd}$ | | 0.6 ± 0.3$^{bcde}$ | | 86.8 ± 28.1$^{def}$ | |
| **AMES_31012_S** | C | 22 ± 4.1$^{ab}$ | 0.02* | 4.1 ± 0.9$^{ab}$ | 0.23 | 10.3 ± 5$^{a}$ | 0.01* | 1.3 ± 0.5$^{a}$ | 0.04* | 170.1 ± 31.7$^{ab}$ | 0.01* |
| | WL | 16.2 ± 2.5$^{cd}$ | | 3.4 ± 0.9$^{bc}$ | | 2.9 ± 1.1$^{bcd}$ | | 0.5 ± 0.2$^{cdef}$ | | 117.7 ± 23.9$^{bcd}$ | |
| **ISU_002_S** | C | 27.3 ± 4.7$^{a}$ | 0* | 4.6 ± 0.9$^{a}$ | 0* | 6.5 ± 1$^{b}$ | 0.16 | 1 ± 0.1$^{ab}$ | 0.11 | 220.2 ± 65.4$^{a}$ | 0.11 |
| | WL | 16.4 ± 3.8$^{cd}$ | | 2.9 ± 0.6$^{cd}$ | | 4.6 ± 2.7$^{bc}$ | | 0.7 ± 0.3$^{bcd}$ | | 162.5 ± 45.1$^{ab}$ | |
| **ISU_182_S** | C | 11.6 ± 1.5$^{de}$ | 0* | 1.2 ± 0.1$^{fg}$ | 0* | 1.9 ± 0.8$^{cd}$ | 0.25 | 0.3 ± 0.2$^{def}$ | 0.33 | 55.8 ± 7.9$^{ef}$ | 0.03* |
| | WL | 7.3 ± 1.2$^{ef}$ | | 0.9 ± 0.1$^{g}$ | | 1.4 ± 0.6$^{cd}$ | | 0.2 ± 0.1$^{ef}$ | | 41.2 ± 11.7$^{ef}$ | |

[A]A "T" or "S" after the accession name denotes if it was predicted to be tolerant or susceptible to waterlogging based on rosette-stage waterlogging experimental data. P-values were determined by Welch's t-test between control and waterlogged replicates. Groups with the same letter next to the standard deviation are not statistically different based on Tukey's HSD test following a two-way ANOVA between all accessions and treatments.
[B]C = control and WL = waterlogged
[C]FW = fresh weight
[D]P-values < 0.05 denoted by asterisk
[E]DW = dry weight

**Supplemental Table 3.** Analysis of variance for biomass traits immediately after waterlogging at the reproductive stage in six natural accessions.

| Trait | Source of Variation | Degrees of Freedom | Sum of Squares | Mean Squares | F value | P value | Signif.[A] |
|---|---|---|---|---|---|---|---|
| Shoot fresh weight (g) | Treatment | 1 | 606.6 | 606.6 | 81.61 | 8.65E-13 | *** |
| | Accession | 5 | 2442.5 | 488.5 | 65.72 | < 2E-16 | *** |
| | Treatment by Accession | 5 | 151.7 | 30.3 | 4.08 | 0.00295 | ** |
| | Residuals | 60 | 446 | 7.4 | | | |
| | Total | 71 | 3646.8 | 1132.8 | | | |
| Shoot dry weight (g) | Treatment | 1 | 5.5 | 5.5 | 18.73 | 5.79E-05 | *** |
| | Accession | 5 | 100.62 | 20.12 | 68.57 | < 2E-16 | *** |
| | Treatment by Accession | 5 | 5.27 | 1.05 | 3.59 | 0.00662 | ** |
| | Residuals | 60 | 17.61 | 0.29 | | | |
| | Total | 71 | 129 | 26.96 | | | |
| Root fresh weight (g) | Treatment | 1 | 89.3 | 89.33 | 25.77 | 3.99E-06 | *** |
| | Accession | 5 | 350.9 | 70.19 | 20.25 | 8.86E-12 | *** |
| | Treatment by Accession | 5 | 107.5 | 21.5 | 6.2 | 0.00011 | *** |
| | Residuals | 60 | 208 | 3.47 | | | |
| | Total | 71 | 755.7 | 184.49 | | | |
| Root dry weight (g) | Treatment | 1 | 1.47 | 1.47 | 25.2 | 4.92E-06 | *** |
| | Accession | 5 | 5.66 | 1.13 | 19.42 | 1.90E-11 | *** |
| | Treatment by Accession | 5 | 1.06 | 0.21 | 3.64 | 0.00614 | ** |
| | Residuals | 60 | 3.5 | 0.06 | | | |
| | Total | 71 | 11.69 | 2.87 | | | |
| Leaf number | Treatment | 1 | 20134 | 20134 | 23.31 | 9.87E-06 | *** |
| | Accession | 5 | 213675 | 42735 | 49.49 | < 2E-16 | *** |
| | Treatment by Accession | 5 | 10764 | 2153 | 2.49 | 0.0407 | * |
| | Residuals | 60 | 51815 | 864 | | | |
| | Total | 71 | 296388 | 65886 | | | |

[A] Level of significance with *= 0.05, **=0.01, ***= 0.001 P-values.

**Supplemental Table 4**. Means and standard deviations of morphological traits collected at the time of maturity after recovery from 1 week of waterlogging at the reproductive stage in eight natural accessions.

| Accession | | Branches | p-val[C] | Height (cm) | p-val | Rep. height (cm) | p-val | Shoot DW[D] (g) | p-val | Maturity | p-val |
|---|---|---|---|---|---|---|---|---|---|---|---|
| **950177_T[A]** | C[B] | 15 ± 4[cdef] | 0.81 | 44.3 ± 3.4[efgh] | 0.57 | 16.9 ± 3.6[efgh] | 0.7 | 1.5 ± 0.5[f] | 0.88 | 47.3 ± 1.5[ab] | 0* |
| | WL | 15.5 ± 4[cdef] | | 43.3 ± 2.5[fgh] | | 15.8 ± 5.5[fgh] | | 1.4 ± 0.2[f] | | 43.2 ± 2[ab] | |
| **AMES_33895_T** | C | 37.8 ± 8.5[a] | 0.02* | 66.6 ± 3.8[bc] | 0.02* | 29.1 ± 3.3[bc] | 0.01* | 4.7 ± 0.7[a] | 0.03* | 32.7 ± 13.3[bcde] | 0.83 |
| | WL | 16.7 ± 10.1[cde] | | 61 ± 3.4[cd] | | 23.2 ± 3.2[cd] | | 3.2 ± 1.2[abcde] | | 34.3 ± 13.3[bcde] | |
| **ISU_077_T** | C | 14.3 ± 6.3[cdef] | 0.01* | 48.4 ± 9.3[efg] | 0.02* | 27.2 ± 8.7[efg] | 0.03* | 2.9 ± 0.3[bcdef] | 0* | 47.8 ± 9.6[ab] | 0* |
| | WL | 3.7 ± 1.5[f] | | 36.1 ± 3.8[h] | | 16 ± 1.8[h] | | 2.2 ± 0.3[def] | | 23.5 ± 2.4[de] | |
| **ISU_378_T** | C | 26 ± 6.5[abc] | 0.05 | 82.2 ± 5[a] | 0.01* | 37 ± 10.2[a] | 0.01* | 3.8 ± 0.6[abc] | 0* | 40.3 ± 9.1[abc] | 0.08 |
| | WL | 16.2 ± 6.1[cdef] | | 72.4 ± 4.5[ab] | | 21.8 ± 3[ab] | | 2.1 ± 0.4[def] | | 32.2 ± 3.6[bcde] | |
| **OSU_5_T** | C | 18.2 ± 6.6[bcde] | 0.03* | 53.6 ± 7.6[def] | 0.84 | 30.3 ± 2.8[def] | 0.2 | 3.6 ± 0.8[abcd] | 0.04* | 45.5 ± 4.6[ab] | 0.26 |
| | WL | 8.5 ± 6.18[ef] | | 54.4 ± 6[de] | | 26.4 ± 5.7[de] | | 2.5 ± 0.7[cdef] | | 39.8 ± 10.3[abcd] | |
| **AMES_31012_S** | C | 17.7 ± 7[cde] | 0.22 | 49.5 ± 6[efg] | 0.02* | 22.3 ± 6.7[efg] | 0.03* | 4.3 ± 1.1[ab] | 0.08 | 31.8 ± 2.5[bcde] | 0* |
| | WL | 11.2 ± 3.9[def] | | 40.9 ± 3.7[gh] | | 14.2 ± 3.1[gh] | | 3.1 ± 1[bcde] | | 22.5 ± 2.7[e] | |
| **ISU_002_S** | C | 30.7 ± 6[ab] | 0* | 52.3 ± 4.9[def] | 0.4 | 23.5 ± 4[def] | 0.01* | 3.5 ± 1.4[abcd] | 0.04* | 42.2 ± 13.6[ab] | 0.03* |
| | WL | 7.5 ± 2.9[ef] | | 49.2 ± 6.9[efg] | | 17.2 ± 3.3[efg] | | 1.9 ± 0.6[ef] | | 24.8 ± 3.1[cde] | |
| **ISU_182_S** | C | 22.8 ± 8.2[bcd] | 0.05 | 70.9 ± 4.1[bc] | 0* | 28.3 ± 6.6[bc] | 0.63 | 3.79 ± 0.68[abc] | 0* | 52.8 ± 10.5[a] | 0.19 |

| | | | | | | |
|---|---|---|---|---|---|---|
| | WL | 12.3 ± 6.7$^{def}$ | 54.3 ± 2.6$^{de}$ | 30.1 ± 6.1$^{de}$ | 1.94 ± 0.37$^{ef}$ | 45.3 ± 7.9$^{ab}$ |

[A] A “T” or “S” after the accession name denotes if it was predicted to be tolerant or susceptible to waterlogging. P-values were determined by Welch’s t-tests between control and waterlogged replicates. Groups with the same letter next to the standard deviation were not statistically different based on Tukey’s HSD test following a two-way ANOVA between all accessions and treatments.

[B] C = control and WL = waterlogged

[C] P-values < 0.05 denoted by asterisk

[D] DW = dry weight

[C] C = control and WL = waterlogged

**Supplemental Table 5.** Means and standard deviations of yield traits and maturity after recovery from 1 week of waterlogging at the reproductive stage in eight natural accessions.

| Accession | | Silicles | p-val[C] | Aborted silicles % | p-val | Seeds per silicle | p-val | Seed weight (g) | p-val |
|---|---|---|---|---|---|---|---|---|---|
| **950177_T[A]** | C[B] | 195.5 ± 37.9[cdef] | 0.14 | 26.1 ± 6.3[a] | 0.01* | 7.3 ± 0.6[abc] | 0.87 | 0.7 ± 0.1[def] | 0.11 |
| | WL | 167 ± 16.7[def] | | 37.4 ± 6.6[a] | | 7.4 ± 1.1[abc] | | 0.6 ± 0.1[defg] | |
| **AMES_33895_T** | C | 522.2 ± 72.9[a] | 0.01* | 22.6 ± 5.6[a] | 0.04* | 7.3 ± 1.2[abc] | 0.69 | 1.8 ± 0.1[a] | 0* |
| | WL | 341.2 ± 121.8[abcd] | | 32.4 ± 8.2[a] | | 7 ± 1.5[abc] | | 0.9 ± 0.1[cde] | |
| **ISU_077_T** | C | 191.3 ± 55.3[cdef] | 0* | 18.4 ± 3.5[a] | 0.21 | 6.8 ± 0.8[abc] | 0.04* | 0.7 ± 0.2[def] | 0* |
| | WL | 55.2 ± 16.8[f] | | 24 ± 9.3[a] | | 5.5 ± 1.2[c] | | 0.2 ± 0.1[g] | |
| **ISU_378_T** | C | 470.8 ± 94.7[ab] | 0* | 28.7 ± 7.2[a] | 0.95 | 7.1 ± 0.9[abc] | 0.56 | 1.1 ± 0.2[bcd] | 0* |
| | WL | 221 ± 45.9[cdef] | | 29 ± 5.2[a] | | 6.8 ± 0.5[abc] | | 0.6 ± 0.1[defg] | |
| **OSU_5_T** | C | 313.3 ± 102.8[bcde] | 0.1 | 35 ± 11.2[a] | 0.35 | 7.2 ± 0.9[abc] | 0.34 | 1 ± 0.3[bcde] | 0.28 |
| | WL | 218 ± 74.7[cdef] | | 29.9 ± 5.6[a] | | 7.8 ± 1.2[ab] | | 0.8 ± 0.2[def] | |
| **AMES_31012_S** | C | 475.5 ± 243.6[ab] | 0.02* | 33.1 ± 23.8[a] | 0.77 | 8.3 ± 1[a] | 0.14 | 1.4 ± 0.5[abc] | 0* |
| | WL | 153 ± 35.8[ef] | | 37.1 ± 21.8[a] | | 7 ± 1.7[abc] | | 0.4 ± 0.2[fg] | |
| **ISU_002_S** | C | 422.3 ± 77.1[ab] | 0* | 20 ± 5.4[a] | 0.11 | 5.7 ± 1[bc] | 0.66 | 1.1 ± 0.3[bcd] | 0* |
| | WL | 120 ± 49.8[f] | | 31.8 ± 14.5[a] | | 5.3 ± 0.6[c] | | 0.2 ± 0.1[g] | |
| **ISU_182_S** | C | 353.5 ± 74[abc] | 0* | 21.8 ± 2.7[a] | 0.01* | 7.2 ± 0.6[abc] | 0.36 | 1.4 ± 0.2[ab] | 0* |
| | WL | 145.5 ± 41.1[ef] | | 31.3 ± 6.1[a] | | 6.7 ± 0.9[abc] | | 0.6 ± 0.1[efg] | |

[A]A “T” or “S” after the accession name denotes if it was predicted to be tolerant or susceptible to waterlogging. Seed weight and shoot dry weight were measured in grams. P-values were determined by Welch’s t-tests between control and waterlogged replicates. Groups with the same letter next to the standard deviation were not statistically different based on Tukey’s HSD test following a two-way ANOVA between all accessions and treatments.
[B]C = control and WL = waterlogged
[C]P-values < 0.05 denoted by asterisk

**Supplemental Table 6.** Analysis of variance for morphology traits at maturity after recovery from 1 week of waterlogging at the reproductive stage in eight natural accessions.

| Trait | Source of Variation | Degrees of Freedom | Sum of Squares | Mean Squares | F value | P value | Signif.[A] |
|---|---|---|---|---|---|---|---|
| Branch number | Treatment | 1 | 3105 | 3105.4 | 78.45 | 1.68E-13 | *** |
| | Accession | 7 | 2562 | 366 | 9.25 | 2.47E-08 | *** |
| | Treatment by Accession | 7 | 1219 | 174.1 | 4.4 | 0.00036 | *** |
| | Residuals | 80 | 3167 | 39.6 | | | |
| | Total | 95 | 10053 | 3685.1 | | | |
| Height (cm) | Treatment | 1 | 1182 | 1182.2 | 43.9 | 3.68E-09 | *** |
| | Accession | 7 | 12408 | 1772.6 | 65.83 | < 2e-16 | *** |
| | Treatment by Accession | 7 | 734 | 104.8 | 3.89 | 0.00107 | ** |
| | Residuals | 80 | 2154 | 26.9 | | | |
| | Total | 95 | 16478 | 3086.5 | | | |
| Reproductive height (cm) | Treatment | 1 | 931.1 | 931.1 | 32.5 | 1.92E-07 | *** |
| | Accession | 7 | 2268.2 | 324 | 11.31 | 7.05E-10 | *** |
| | Treatment by Accession | 7 | 614.9 | 87.8 | 3.07 | 0.00653 | ** |
| | Residuals | 80 | 2292.1 | 28.7 | | | |
| | Total | 95 | 6106.3 | 1371.6 | | | |
| Shoot dry weight (g) | Treatment | 1 | 35.72 | 35.72 | 62.04 | 1.41E-11 | *** |
| | Accession | 7 | 47.43 | 6.78 | 11.77 | 3.33E-10 | *** |
| | Treatment by Accession | 7 | 7.85 | 1.12 | 1.95 | 0.0729 | NS |
| | Residuals | 80 | 46.06 | 0.58 | | | |
| | Total | 95 | 137.06 | 44.2 | | | |
| Maturity (days) | Treatment | 1 | 2100 | 2100 | 31.69 | 2.59E-07 | *** |
| | Accession | 7 | 4361 | 623 | 9.4 | 1.87E-08 | *** |
| | Treatment by Accession | 7 | 1365 | 194.9 | 2.94 | 0.00858 | ** |
| | Residuals | 80 | 5302 | 66.3 | | | |
| | Total | 95 | 13128 | 2984.2 | | | |

[A]Level of significance with *, **, *** indicates P-values at 0.05, 0.01, and 0.001, respectively.

**Supplemental Table 7**. Analysis of variance for yield traits at maturity after recovery from 1 week of waterlogging at the reproductive stage in eight natural accessions.

| Trait | Source of Variation | Degrees of Freedom | Sum of Squares | Mean Squares | F value | P value | Signif.[A] |
|---|---|---|---|---|---|---|---|
| Silicle number | Treatment | 1 | 870585 | 870585 | 108.27 | < 2E-16 | *** |
| | Accession | 7 | 763750 | 109107 | 13.57 | 1.96E-11 | *** |
| | Treatment by Accession | 7 | 216301 | 30900 | 3.84 | 0.00119 | ** |
| | Residuals | 80 | 643293 | 8041 | | | |
| | Total | 95 | 2493929 | 1018633 | | | |
| Aborted silicles % | Treatment | 1 | 829 | 829.5 | 7.21 | 0.00882 | ** |
| | Accession | 7 | 1616 | 230.9 | 2.01 | 0.06437 | NS |
| | Treatment by Accession | 7 | 746 | 106.6 | 0.93 | 0.49112 | NS |
| | Residuals | 80 | 9205 | 115.1 | | | |
| | Total | 95 | 12396 | 1282.1 | | | |
| Seeds per pod | Treatment | 1 | 4.17 | 4.17 | 3.71 | 0.0576 | NS |
| | Accession | 7 | 46.16 | 6.6 | 5.87 | 1.61E-05 | *** |
| | Treatment by Accession | 7 | 9.03 | 1.29 | 1.15 | 0.3417 | NS |
| | Residuals | 80 | 89.85 | 1.12 | | | |
| | Total | 95 | 149.21 | 13.18 | | | |
| Total seed weight (g) | Treatment | 1 | 9.49 | 9.492 | 151.14 | 4.11E-11 | *** |
| | Accession | 7 | 5.75 | 0.822 | 13.09 | < 2E-16 | *** |
| | Treatment by Accession | 7 | 2.54 | 0.363 | 5.78 | 1.93E-05 | *** |
| | Error | 80 | 5.02 | 0.06 | | | |
| | Total | 95 | 22.8 | 10.737 | | | |

[A]Level of significance with *, **, *** indicates P-values at 0.05, 0.01, and 0.001, respectively

**Figures**

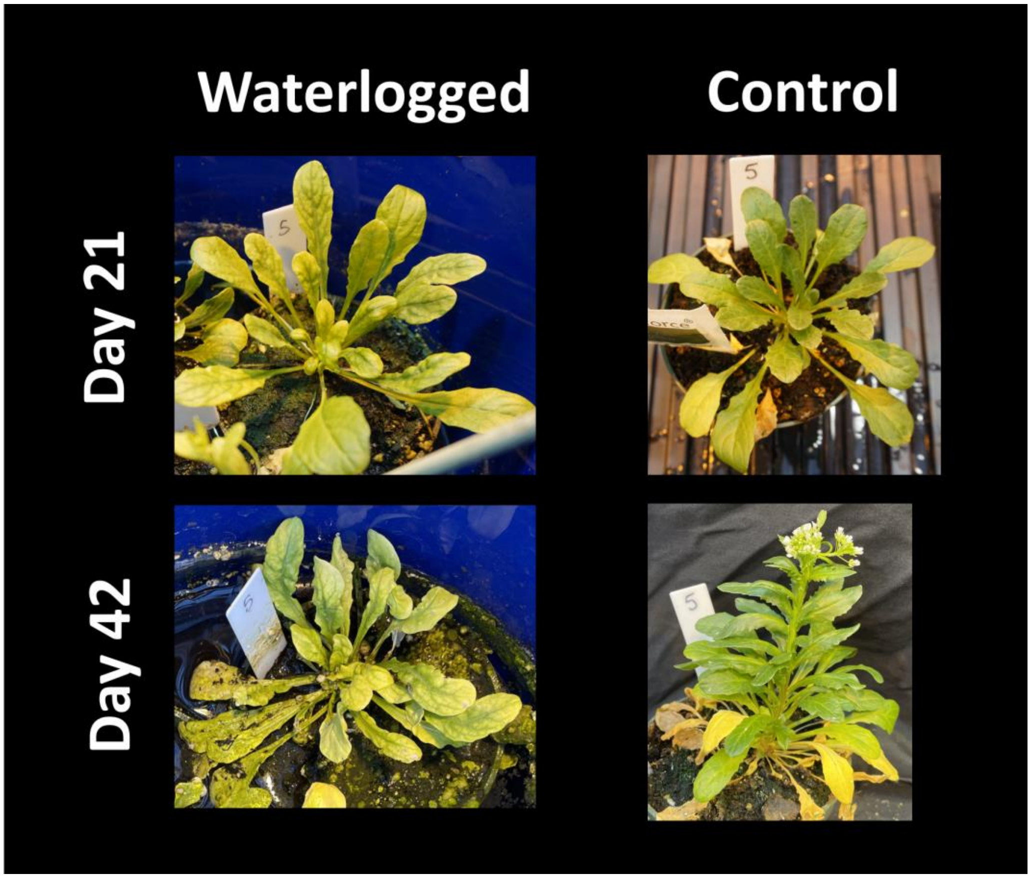


**Supplemental Figure 1.** Pictures of leaf chlorosis observed under rosette-stage waterlogging in AMES_31013.

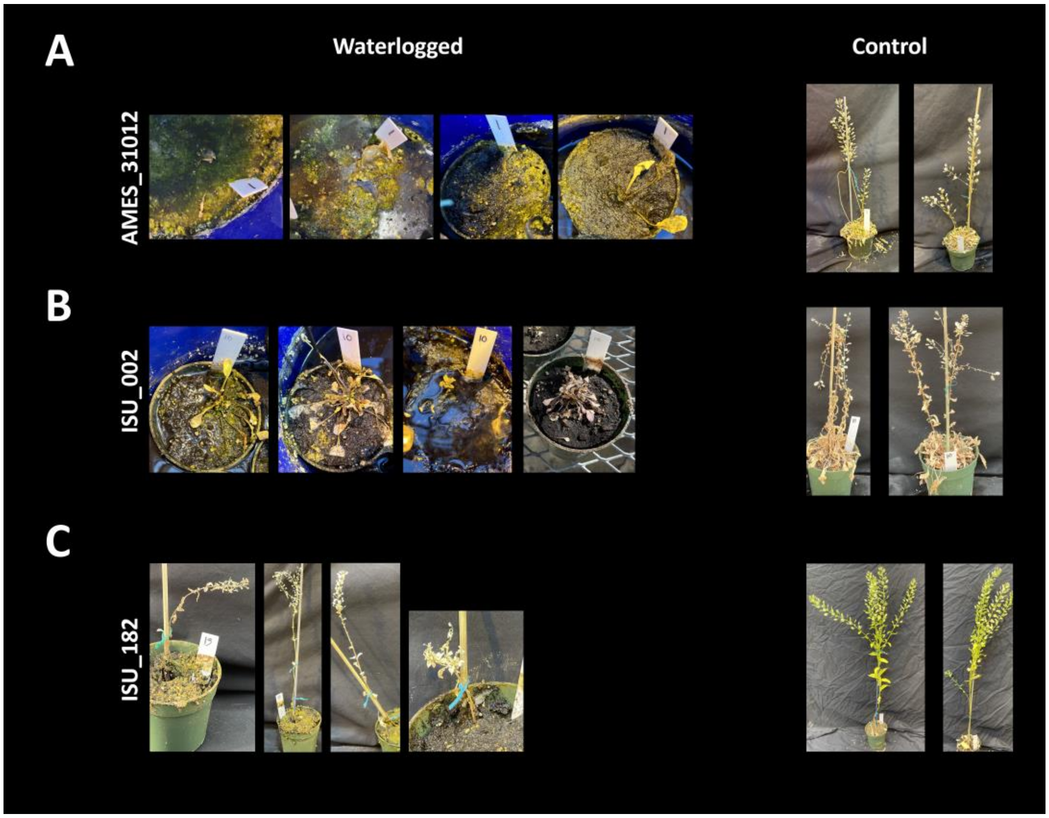


**Supplemental Figure 2.** Pictures of dead waterlogged plants, and their mature control counterparts, for A) AMES_31012, B) ISU_002, and C) ISU_182 after waterlogging at the rosette stage. These 3 accessions were chosen as "susceptible" lines for the reproductive stage experiments.

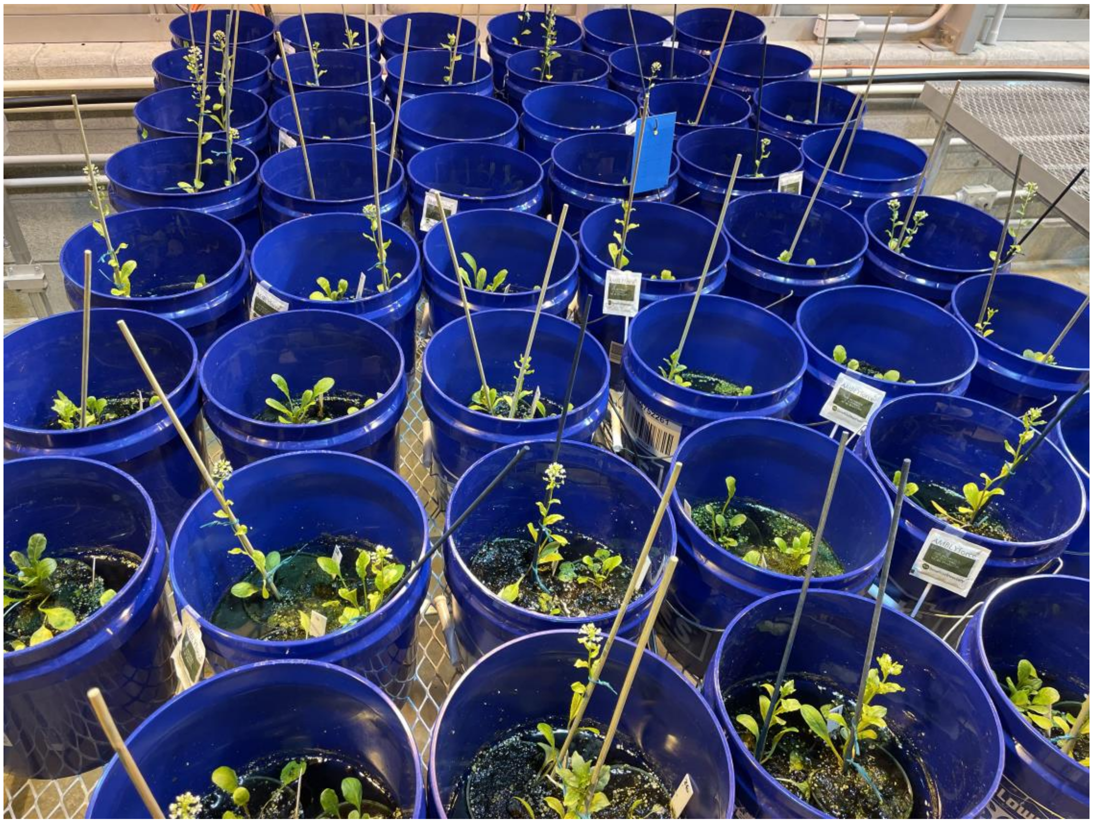

**Supplemental Figure 3.** Picture of pennycress plants after 2 weeks of waterlogging that began at the rosette stage.  About half of the plants were bolting or flowering, while the other half remained in the rosette stage.

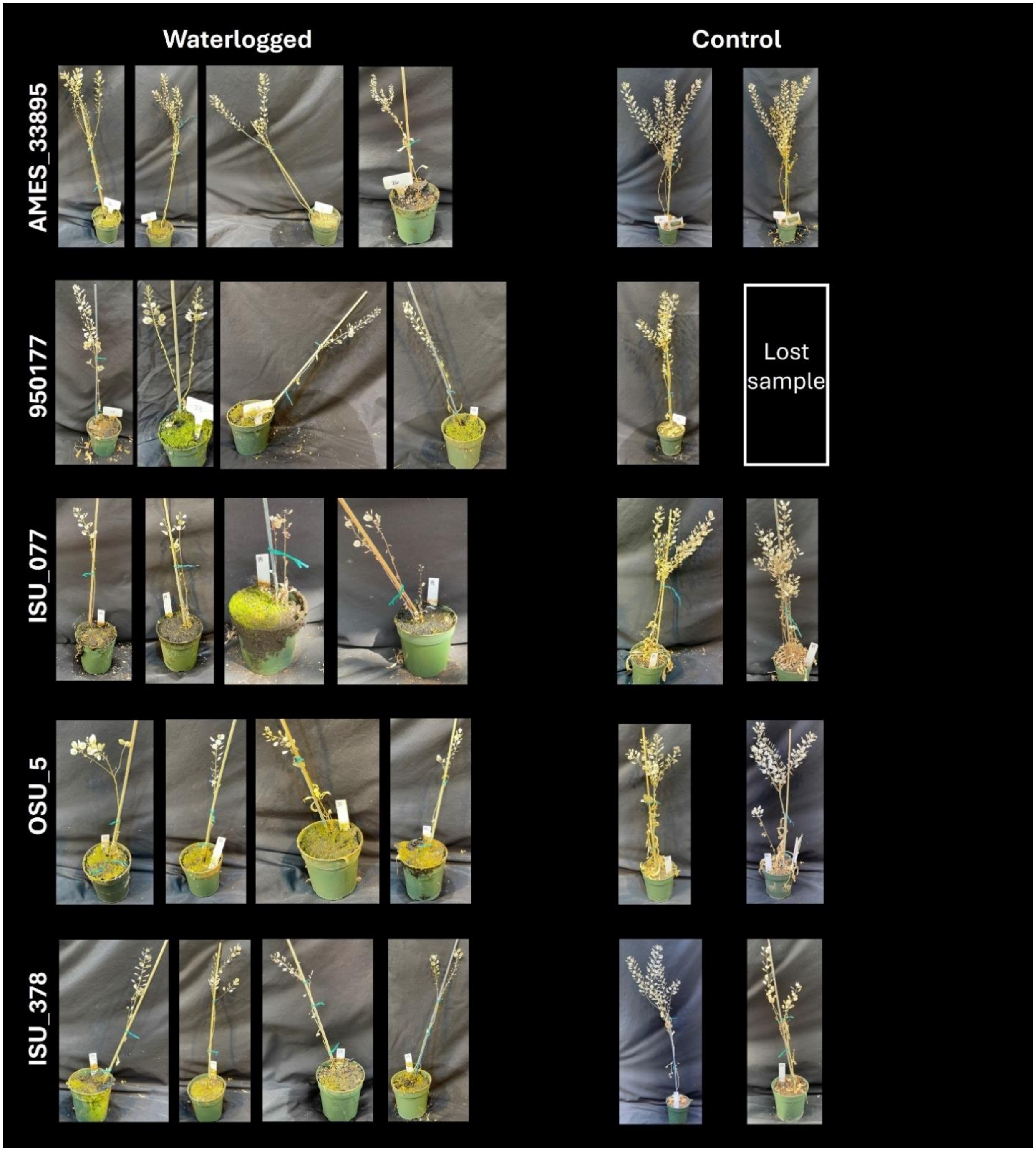


**Supplemental Figure 4.** Pictures of mature waterlogged and control replicates from 5 accessions after continuous waterlogging at the rosette stage. These 5 accessions were chosen as "tolerant" lines for the reproductive stage experiment. One control replicate from 950177 was lost due to aphids.
